# ACE2 mimetic antibody potently neutralizes all SARS-CoV-2 variants and fully protects in XBB.1.5 challenged monkeys

**DOI:** 10.1101/2023.07.18.549530

**Authors:** Craig Fenwick, Priscilla Turelli, Yoan Duhoo, Kelvin Lau, Cécile Herate, Romain Marlin, Myriam Lamrayah, Jérémy Campos, Line Esteves-Leuenberger, Alex Farina, Charlène Raclot, Vanessa Genet, Flurin Fiscalini, Julien Cesborn, Laurent Perez, Nathalie Dereuddre-Bosquet, Vanessa Contreras, Kyllian Lheureux, Francis Relouzat, Rana Abdelnabi, Caroline S. Foo, Johan Neyts, Pieter Leyssen, Yves Lévy, Florence Pojer, Henning Stahlberg, Roger Le Grand, Didier Trono, Giuseppe Pantaleo

**Affiliations:** Service of Immunology and Allergy, Department of Medicine, Lausanne University Hospital and University of Lausanne, Lausanne, Switzerland; School of Life Sciences, Ecole Polytechnique Fédérale de Lausanne, Lausanne, Switzerland; School of Basic Sciences, Ecole Polytechnique Fédérale de Lausanne and Faculty of Biology and Medicine, UNIL, Lausanne, Switzerland; CEA, Université Paris Sud 11, INSERM U1184, Center for Immunology of Viral Infections and Autoimmune Diseases, IDMIT Department, IBFJ, Fontenay-aux-Roses, France; KU Leuven Department of Microbiology, Immunology and Transplantation, Rega Institute for Medical Research, Laboratory of Virology and Chemotherapy, B-3000 Leuven, Belgium; VRI, Université Paris-Est Créteil, Faculté de Médicine, INSERM U955, 94010 Créteil, France; Inserm U955, Equipe 16, Créteil, France; AP-HP, Hôpital Henri-Mondor Albert-Chenevier, Service d’Immunologie Clinique et Maladies Infectieuses, Créteil, France; Swiss Vaccine Research Institute, Lausanne University Hospital and University of Lausanne, Switzerland

**Keywords:** SARS-CoV-2, neutralizing antibodies, Variants of concern, Omicron

## Abstract

The rapid evolution of SARS-CoV-2 to variants with improved transmission efficiency and reduced sensitivity to vaccine-induced humoral immunity has abolished the protective effect of licensed therapeutic human monoclonal antibodies (mAbs). To fill this unmet medical need and protect vulnerable patient populations, we isolated the P4J15 mAb from a previously infected, vaccinated donor, with <20 ng/ml neutralizing activity against all Omicron variants including the latest XBB.2.3 and EG.1 sub-lineages. Structural studies of P4J15 in complex with Omicron XBB.1 Spike show that the P4J15 epitope shares ∼93% of its buried surface area with the ACE2 contact region, consistent with an ACE2 mimetic antibody. Although SARS-CoV-2 mutants escaping neutralization by P4J15 were selected *in vitro*, these displayed lower infectivity, poor binding to ACE2, and the corresponding ‘escape’ mutations are accordingly rare in public sequence databases. Using a SARS-CoV-2 XBB.1.5 monkey challenge model, we show that P4J15 confers complete prophylactic protection. We conclude that the P4J15 mAb has potential as a broad-spectrum anti-SARS-CoV-2 drug.

## MAIN

The SARS-CoV-2 virus which lead to the global COVID-19 pandemic is responsible for >767 million confirmed infections and almost 7 million fatalities worldwide (WHO Coronavirus (COVID-19) Dashboard https://covid19.who.int/) (*1*). Enormous efforts by the scientific and medical communities in vaccine, antiviral drugs and monoclonal antibody development have allowed most people to return to normal lives after the peak of the pandemic typified in most parts of the world by lockdowns, isolation, and overwhelmed health care networks. However, these hard-fought victories are being eroded by the continued regional and international spread of variants of concern (VOC), which are both more transmissible and more resistant to immune responses (*2–4*). SARS-CoV-2 infection leading to COVID-19 disease is of increased concern due to the rapidly waning immunity in the general population, the apathy that has developed for receiving vaccine boosts and the reduced levels of neutralizing antibodies generated by even the most recent bivalent vaccines (which include the ancestral SAR-CoV-2 and Omicron BA.1 or BA.4/5 Spike) against current VOCs (*5*). Alarmingly, these factors contribute to SARS-CoV- 2 infection being a leading cause of death in children and adolescents up to 19 years of age, accounting for 2% of all deaths in this age group in the year prior to August 2022 (*6*). The greatest unmet medical need is the >30 million immunocompromised individual in the US and Europe alone that are at high risk of infection due to their inability to mount a protective humoral immune response following vaccination. These at-risk individuals include people with blood and immune cancers, transplantation patients and recipients of immunosuppressive drugs, all of which account for >40% of hospitalizations with breakthrough SARS-CoV-2 infections. Since the emergence of the BQ.1, BQ.1.1 and XBB.1 lineages in the fall of 2022, all authorized therapeutic mAbs have become almost completely ineffective, including Evusheld, the combination of tixagevimab (AZD8895) and cilgavimab (AZD1061) mAbs that was designed for therapeutic and prophylactic purposes (*7*). Although there are recent reports in the literature identifying mAbs with some breadth of neutralizing activity, most are significantly less potent against the circulating variants than antibodies that previously demonstrated protection in the clinic (*8–11*). Furthermore, the SARS-CoV-2 virus will continue to evolve due to both the tremendous pool of circulating viruses and selective pressures exerted by immune responses present in the general population, necessitating that new mAbs be developed to counter currently circulating variants and anticipate future adaptations. Here we report the isolation of the human mAb, P4J15, that binds the Spike receptor binding domain (RBD) and through blocking ACE2 receptor binding, exerts a potent neutralizing activity against all current SARS-CoV-2 variants. Structural studies reveal that this broad activity can be attributed to the high binding surface area of the P4J15 epitope, which includes residues essential for efficient ACE2 binding and infection. Live virus resistance studies confirmed that P4J15 escape mutants selected in cell culture were poorly infectious, owing to RBD mutations strongly reducing affinity for the ACE2 receptor. Accordingly, the corresponding escape mutations were found only at low frequency in the GISAID sequence database, confirming their detrimental effect on virus fitness. Finally, P4J15 conferred near complete protection from infection in hamster and monkey live virus challenge studies performed with Omicron BA.5 and XBB.1.5 variants, respectively. With this demonstrated *in vivo* efficacy against the most recent variants, neutralizing potency and breadth of protection, we propose that P4J15 could be a strong candidate for clinical development.

## RESULTS

### P4J15 is a potent and broadly neutralizing anti-SARS-CoV-2 monoclonal antibody

As part of a longitudinal study to monitor the waning humoral immune response in a cohort of 20 donors, we performed routine serum testing of anti-Spike and anti-Nucleocapsid antibodies. One donor with weak neutralizing antibody levels consistent with prior SARS-CoV-2 infection, received two doses of the mRNA-1273 vaccine in late 2021 and became SARS-CoV-2 positive four weeks later during the Omicron BA.1 wave. Two months later, this donor had among the highest serum antibody levels in all donors tested, with excellent breadth of neutralization against Omicron BA.1 and a panel of other pre-and post-Omicron SARS-CoV-2 variants in a trimeric Spike-ACE2 surrogate neutralization assay (*12*). To focus our mAb screening efforts, we sorted Omicron BA.1 Spike-binding memory B cells and identified a panel of 16 Spike binding antibodies by profiling antibody supernatants from B cell clones. The P4J15 mAb, produced by expression of paired heavy and light chains in ExpiCHO cells, showed the highest affinity for the ancestral, Alpha, Beta, Gamma, Delta, Omicron BA.1, BA.1.1, BA.2, BA.2.75.2, BQ.1, BQ.1.1, XBB.1 and RaTG13 Spike trimer, while showing only low levels of binding to SARS-CoV-1 Spike (**Supplementary Data Fig. 1a-b**). Profiling studies in a Luminex Spike binding assay showed that the purified P4J15 mAbs bound SARS-CoV-2 Spike proteins in our panel with IC50 values ranging from 0.6 to 1.5 ng/ml (**Supplementary Data Fig. 1b**) and RaTG13 Spike with an IC50 of 11 ng/ml. Using our Spike-ACE2 surrogate neutralization assay, we also found that P4J15 had the most potent and broadest activity in blocking ACE2-binding to Spike trimers from our extensive panel of SARS-CoV-2 variants, with IC50 values below 5 ng/ml (**Supplementary Data Fig. 2a-b**). These studies indicate that P4J15 has a superior affinity in binding and Spike-ACE2 blocking profile compared to a panel of approved or clinically advanced anti-SARS-CoV-2 mAbs including AZD8895 and AZD1061 from AstraZeneca (*13*), ADG-2 from Adagio (*14*), bebtelovimab from Eli Lilly (*15*) and S309/sotrovimab from Vir/GSK(*16*).

Cross-competitive Spike trimer binding studies performed with our panel of comparator anti-SARS-CoV-2 mAbs and P2G3/P5C3 mAbs previously described by our group (*17, 18*) revealed that P4J15 binds an overlapping epitope with AZD8895 and P5C3 class 1 mAbs, although neither of these mAbs identified early in the pandemic significantly binds to post-BA.4/5 Omicron variants. No cross-competitive binding was observed between P4J15 and the class 3 mAbs sotrovimab, bebtelovimab and P2G3, whereas the class 4 mAb ADG-2 showed mixed competition results with P4J15, depending on which antibody was bound to Spike first. (**Supplementary Data Fig. 3**).

### P4J15 outperforms clinically relevant antibodies in pseudovirus-based neutralization assays

We next evaluated the activity of P4J15 compared with a panel of clinically approved mAbs in pseudotyped lentiviral and SARS-CoV-2 virus-like particle (VLP) cell-based neutralization assays. P4J15 demonstrated potent neutralizing activity against lentiviruses pseudotyped with Spike from the ancestral 2019-nCoV (D614G), Alpha, Beta, Gamma, Delta, Omicron BA.1, BA.4/5 and BA.2.75.2 VOCs with EC50 values of 41, 14, 19, 16, 9, 5 and 9 ng/ml, respectively (**Fig. 1a-b**). In contrast to all other reference antibodies, P4J15 strongly neutralized the Omicron BQ.1, BQ.1.1, XBB.1 Spike pseudoviruses with an IC50 values of 6, 9 and 14 ng/ml, respectively (**Fig. 1b-c**) and showed no significant loss of activity as compared to the other VOCs. In parallel testing, P4J15 was 13- to 300-fold more potent than sotrovimab, while the Evusheld dual combination of neutralizing mAbs, and bebtelovimab were almost inactive against BQ.1.1 and XBB.1 VOCs with EC50s >8700 ng/ml. As Spike is incorporated at the plasma membrane in pseudotyped lentivectors and in the ER-Golgi intermediate compartment (ERGIC) in SARS-CoV-2 viruses, we decided to use a SARS-CoV-2-based particle (VLPs) as a second system to profile P4J15 efficacy. These VLP-based assays confirmed results obtained in the lentiviral assay pseudotyped with Spike from Delta BA.1, BA.4/5, BA.2.75.2, BQ.1, BQ.1.1 and XBB.1 (EC50 values of 15, 2, 4, 10, 10, 13 and 13 ng/ml, respectively) (**Fig. 1b** and **1d**). VLP neutralizing assays also revealed that P4J15 retained full neutralizing activity against the latest circulating Omicron sublineages XBB.1.5, CH.1.1, XBB.1.16, XBB.1.16.1, XBB.2.3 and EG.1 with EC50 values of 6, 10, 9, 13 18, and 16 ng/ml (**Fig. 2e** with results summarized in **Fig. 2b**).

**Figure 1:**
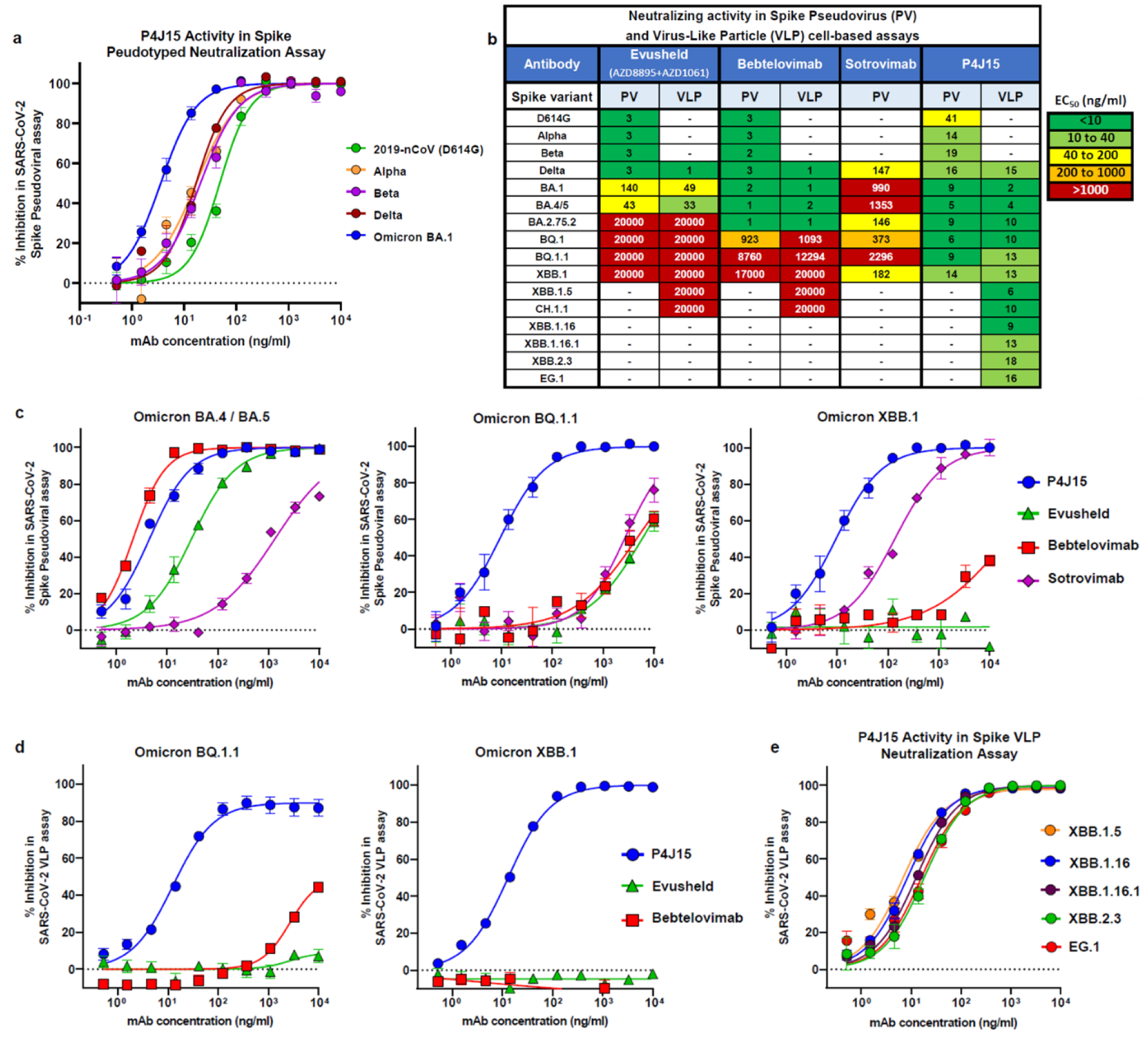
P4J15 exhibits potent and broad neutralizing activity against spike-coated pseudoviruses. **a**) Neutralization of lentiviral particles pseudotyped with SARS-CoV-2 Spike expressing the ancestral 2019-nCoV (D614G), Alpha, Beta, Delta or Omicron BA.1 variants of concern in HEK293T ACE2/TMPRSS2 cell infection assays. Replicates in concentration response curves were n=6 for all Spike pseudoviruses. **b**) Heatmap table showing IC50 neutralization potencies for P4J15 and reference antibodies Evusheld (combination of AZD8895 and AZD1061), bebtelovimab and sotrovimab evaluated in spike-coated pseudovirus and SARS-CoV-2 virus-like particle cell-based assays. **c**) Concentration response inhibition curves for Omicron BA.4/BA.5, BQ.1.1 and XBB.1 Spike pseudotyped lentivirus cell-based neutralization assays (n=6). **d**) Concentration response inhibition curves for Omicron BQ.1.1 and XBB.1 Spike pseudotyped SARS-CoV-2 VLP cell-based neutralization assays (n=8 for Evusheld and n=12 for remaining mAbs). **e**) Concentration response inhibition curves for P4J15 in XBB.1.5, XBB.1.16, XBB.1.16.1, XBB.2.3 and EG.1 Spike pseudotyped VLP cell-based neutralization assays (n=4). Mean values ± SEM are shown.

**Figure 2.**
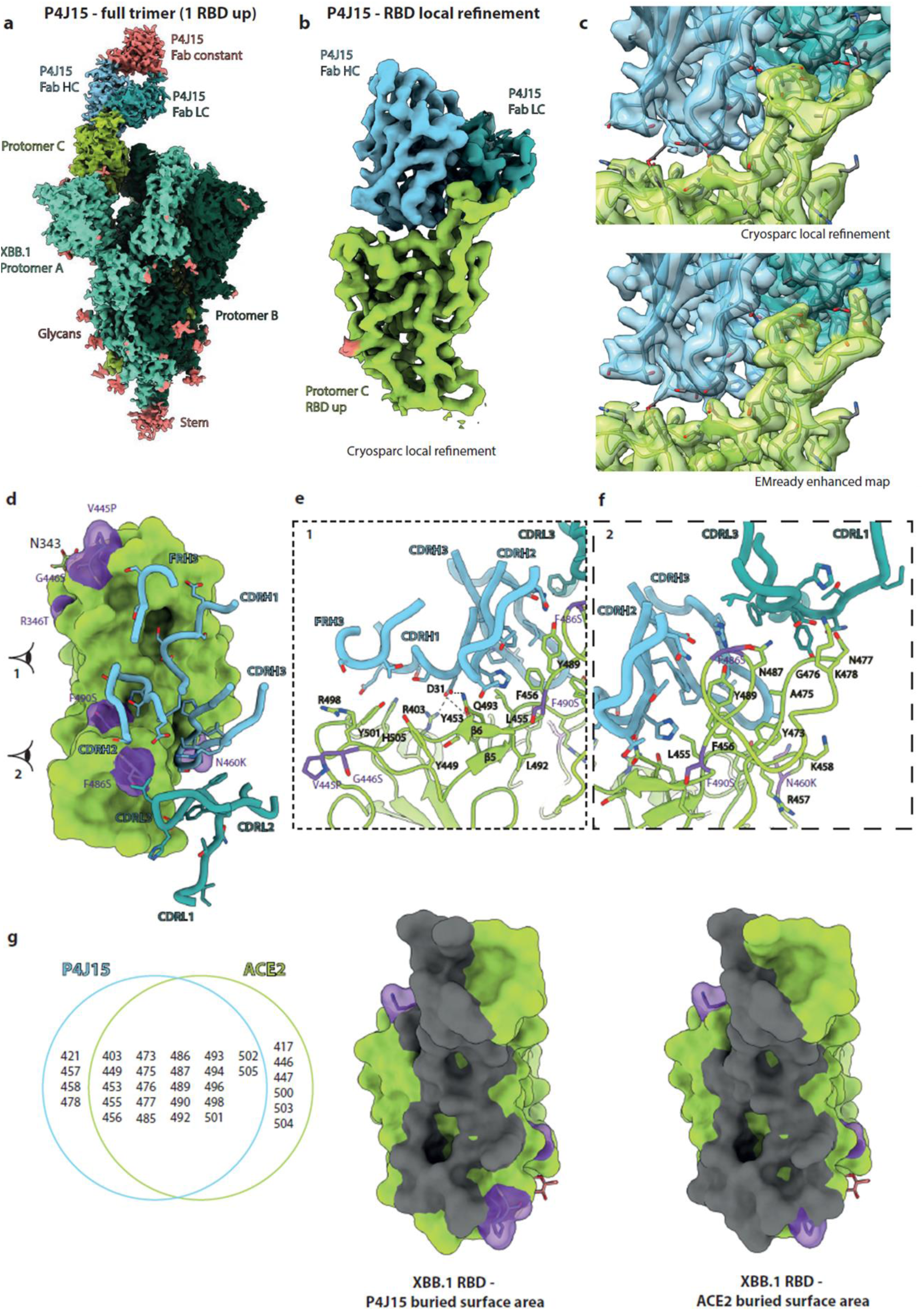
P4J15 binds the full-length Omicron Spike. **a**) Cryo-EM composite density map of the full-length Omicron XBB.1 Spike bound to one P4J15 Fab fragment. Spike protomers are colored in light green, green and dark green, while P4J15 heavy chain variable region is cyan, the light chain variable region turquoise and Fab constant regions pink. **b**) Cryosparc local refinement map of RBD in the up configuration (light green) bound by P4J15 heavy and light chains. **c**) Zoomed-in view of RBD – P4J15 interaction with ribbon structure representation of both and semi-transparent surface representations of cryosparc local refinement and EMReady enhanced maps, shown in top and bottom panels, respectively. **d**) Top view representation of the RBD in green and P4J15 heavy and light chain contact loops shown in cyan and turquoise, respectively. **e)-f)** Front view of the RBD interaction region with P4J15 as indicated by the eye 1 and eye 2 as shown in panel d) for e) and f), respectively. RBD is represented as a ribbon structure in green, while P4J15 heavy chain CDRs and frame region 3 (FRH3) are shaded cyan and light chain CDRs are shaded turquoise. Key RBD and P4J15 contact residues are labelled and represented in stick format. **g**) Venn diagram showing common contact residues on RBD shared between P4J15 and ACE2. Buried surface area for P4J15 and ACE2 are shaded dark grey on the green space filled representation of the RBD. In panels d) and g), the XBB.1 mutations relative to Omicron BA.4 are shaded purple.

### Cryo–electron microscopy structure of P4J15 Fab in complex with the Spike trimer

To understand the structural basis of P4J15 potent neutralization of SARS-CoV-2 variants of concern, the complex formed by Omicron XBB.1 SARS-CoV-2 Spike trimer and P4J15 Fab fragments was characterized using single particle cryo-electron microscopy (Cryo-EM). The single particle cryo-EM reconstruction of the Omicron XBB.1 Spike trimeric ectodomain bound to the Fab at a 3.01 Å resolution and the P4J15 Fab binding RBD in the up-or open-conformation (**Fig. 2a** and **Supplementary data Fig. 4a-e** and **6**). P4J15 binds RBD with a buried surface area of approximately 828 Å^2^ as a Class 1 neutralizing mAb, recognizing an epitope on the SARS-CoV-2 RBD that largely overlaps with the ACE2 receptor binding motif. To characterize the P4J15 paratope and epitope interface in detail, we performed local refinement of the P4J15 Fab-RBD interacting region and reached a resolution of 3.85 Å with well-defined density, allowing clear interpretation of sidechain positions at the interface. We also used EMReady (*19*), a deep learning tool, to enhance the quality even further (**Fig. 2b-c** and **Supplementary data Fig. 4**-**5**). The P4J15 paratope interactions are mediated mainly through electrostatic and hydrophobic contacts and involve 26 residues of the RBD, bound by the three heavy chain CDRs, two light chain CDRs and residues of the heavy chain Frame region 3 (FRH3) of the P4J15 mAb. The CDRH1, CDRH2 and FRH3 loops of P4J15 (**Fig. 2d-f** and **Supplementary data Fig. 7**-**8**) form contacts with the Gly447-Phe456 and Gly485- His505 saddle-like region of the RBD that encompasses the β5 and β6 antiparallel beta sheet of the ACE2 binding region (**Fig. 2e**). CDRH3 sits upon the RBD Arg454-Lys458 loop and together with the CDRL1 and CDRL3 forms additional contacts with the Tyr473-Lys478 and Ser486-Tyr489 of the ACE2 interface region (**Fig. 2f**). These contact residues are further illustrated in **Figure 2g** with a structural model of the RBD viewed from above and the P4J15 contact buried surface on the RBD shaded in dark grey and compared to the ACE2 contact residues. It is important to underscore that 22 of these P4J15 contact residues on the RBD are shared with key contacts formed between the RBD (**Fig. 2g**) and the ACE2 receptor, which is essential for virus binding and infection of target cells. The common area on the RBD for these 22 residues that contact P4J15 and ACE2 is on average 774 Å^2^, which is almost 93% of the P4J15 epitope and 87% of the 887 Å^2^ contact area with ACE2. Based on these observations, we propose that P4J15 may act as an ACE2 mimetic antibody and that it will be difficult for the virus to develop resistance mutations that completely disrupt the P4J15-RBD interaction without adversely affecting the ACE2-RBD interaction.

### P4J15 viral escape mutations have reduced infectivity and ACE2 binding

To gain insight into the predicted clinical value of P4J15 and variant residues in the SARS-CoV-2 Spike that may affect the mAbs neutralizing activity, we characterized the emergence of escape mutants in live virus tissue culture studies. For this, we grew SARS-CoV-2 Omicron BA.2.75.2 and Omicron BQ.1 variants in the presence of sub-optimal neutralizing doses of P4J15 for three passages to generate a heterogeneous viral population, before switching to stringent mAb concentrations in order to select authentic escapees (**Fig. 3a**). Viral genome sequencing of these mAb-resistant mutants pointed to the importance of Spike substitutions F456S, A475D, G476D, N477D/K, N487S/D/K escaping P4J15 in the BA.2.75.2 selection studies and G476D, N487S/T/D/K, Y489H substitutions identified with BQ.1 virus studies (**Fig. 3b**). The identified mutations were then generated by site-directed mutagenesis in a Spike BA.2.75.2 and Spike BQ.1 expression vectors and used to generate mutant Spike virus-like particles (VLPs), allowing us to measure the impact of these mutations on the neutralizing capacity of P4J15 and on viral infectivity. Spike mutations that conferred a near complete loss of neutralizing activity in the BA.2.75.2 VLP assay were F456S, A475D, G476D, N487D, N487K, and N487T while N477D, N477K and N487S conferred a 14- to 29-fold loss of activity (**Fig. 3c**). Similarly, in the BQ.1 Spike VLP assay, G476D, N487D, N487K, and N487T induced resistance to neutralization by P4J15 along with the Y489H mutation, while the N487S mutation conferred only partial resistance (**Fig. 3c**). However, the infectivity of select VLPs was reduced with Spike proteins harboring many of the escapee mutations (**Fig. 3d**) including A475D and N487D in the BA.2.75.2 Spike and N487D, N487K and N487T in BQ.1 Spike. Furthermore, using the ACE2-RBD interactive tool developed by Jesse Bloom’s lab(*20*), it was found that all the Spike escape mutations in RBD induced a strong, 1- to 3-log reduction in binding affinity for ACE2 relative to the Omicron BA.2 Spike reference strain (**Fig. 3e**). The reduced binding affinity of ACE2 for the Spike RBD escape mutations was expected based on our structural data in **Figure 2e-g**, as these residues are important for both P4J15 and ACE2 binding to the RBD. Furthermore, we examined the GISAID EpiCoV database to determine the frequency of the Spike mutations mediating escape to P4J15 neutralization. They were all found to be extremely rare and present in less than 0.0051% of the >15’6 million sequences deposited as of June 2023 (**Fig 3c**). Therefore, escape mutations to P4J15 are only present at very low frequency in viruses isolated from infected individuals, consistent with the marked reduction in infectivity and/or ACE2 binding measured *in vitro* for the corresponding viruses and Spike proteins (**Fig. 3d-e**). Finally, to confirm the barrier to P4J15 viral resistance imposed by the obligatory Spike-ACE2 interaction, we bioinformatically identified the rare but most common amino acid substitutions at positions identified at the P4J15-RBD contact site and in our resistance studies. As shown in **Figure 3f**, with substitutions made to the Omicron BA.5 or BQ.1 Spike, none of the mutations tested, including N417D, V445D, G446D, N450D, L455F, F456L, K458H, S459P, A475V, G476S, N477G, T478R, G485D, P491S, S494P and G504D that are within or adjacent to the P4J15-RBD contact site, significantly reduced the neutralizing potency of P4J15. These studies support the hypothesis that the large, buried surface area bound by P4J15 translates into the antibody’s ability to lose some of these contacts without affecting the overall binding and neutralizing properties.

**Figure 3:**
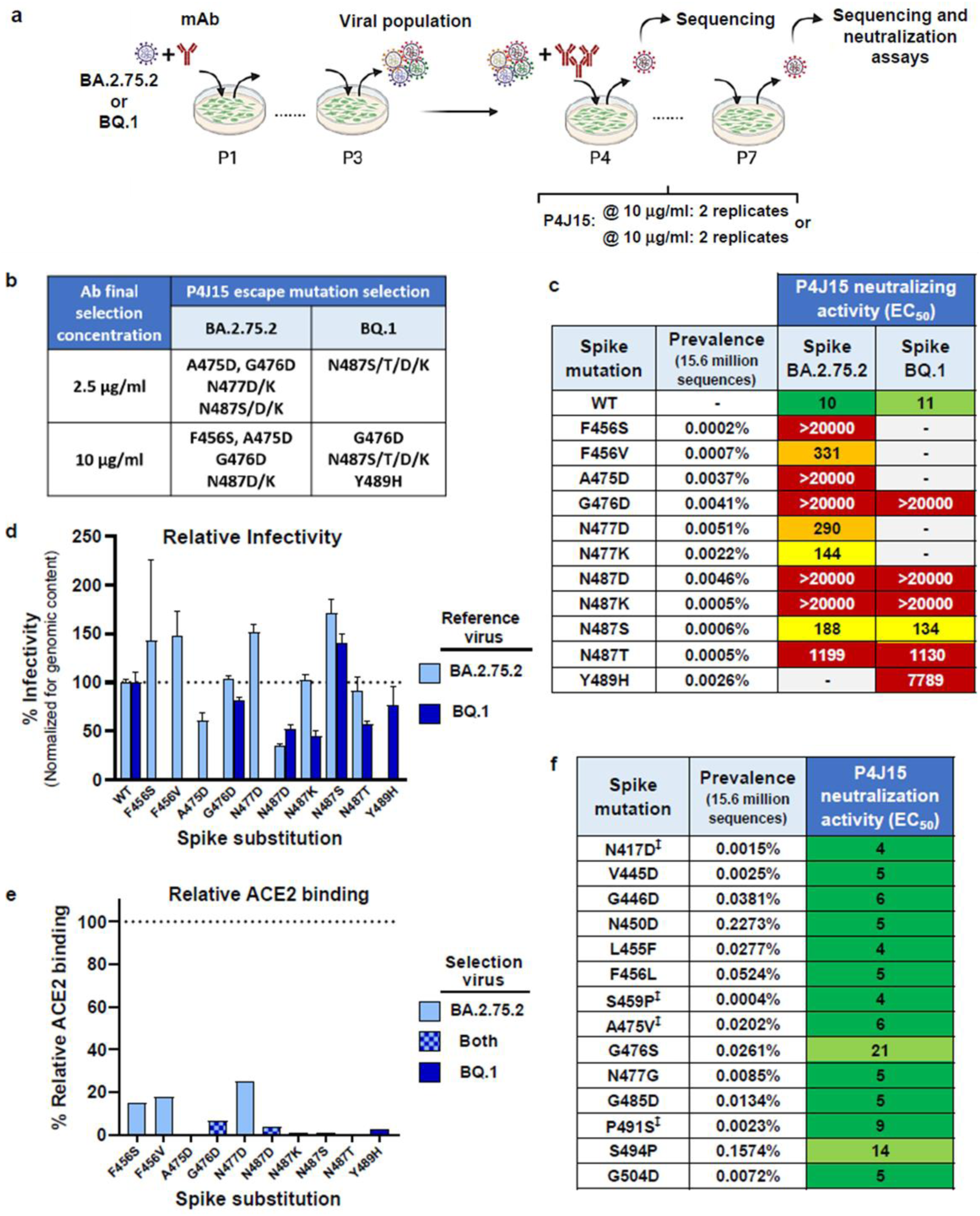
Identification and characterization of escape mutations to P4J15. **a)** Schematic representation of escapees selection. Omicron BA.2.75.2 and BQ.1 replicative isolates were used to infect VeroE6 cells (MOI of 0.2) each in duplicates in presence of suboptimal concentrations of antibodies. Supernatants were collected, diluted 40-fold and used to infect cells for two more passages in the same conditions (P1 to P3). Putative viral escapees were further selected by serial passages of 2-fold diluted supernatants pre-incubated with high concentrations of antibodies (two concentrations, each tested in duplicates). Viral RNA extracted from supernatants collected at passage 5 was deep-sequenced. **b)** Mutations identified across escape selection experiments are indicated in the table. **c)** Prevalence in GISAID sequence database is indicated for each identified mutation. BA.2.75.2 or BQ.1 Spikes were mutated accordingly with the identified residues and pseudotyped VLPs produced and tested in conventional neutralization assays. Heatmap table overviews the EC50 value neutralizing potency of P4J15 against the different VLPs. **d**) Infectivity of the VLPs pseudotyped with the different Spike mutations is shown relative to the parent VLP produced with either BA.2.75.2 or BA.1 Spike. Transduction efficiency was monitored by Luciferase activity in the VLPs transduced HEK293T ACE2/TMPRSS2 cells (n= 16 for all except for G476D and N487 mutations where each have been tested in n=8 replicates). Infectivity is given for the same amount of each infectious VLP as determined by genome content of the stocks. Mean values ± SD are shown, and Kruskal-Wallis test shows significantly reduced infectivity of A475D, N487D (p=0.0065 and p<0.0001, respectively) for BA.2.75.2 VLPs and N487D, N487K, N487T (each p<0.0001) for BQ.1 VLPs. **e)** Relative binding of ACE2 to RBD with the indicated amino acid substitutions as determined using the ACE2-RBD interactive tool developed by Jesse Bloom’s laboratory with Omicron BA.2 used as the reference variant. **f)** Pseudotyped lentiviruses or VLPs produced with rare but most common amino acid substitutions at positions identified at or near the P4J15-RBD contact site and in our resistance studies. Mutations were incorporated in the Omicron BA.4 /BA.5 Spike for lentiviruses and BQ.1 Spike for VLPs (indicated by ‡) with the prevalence in the GISAID sequence database indicated of each variant substitution. Heatmap tables (with same color ranges as in Figure 1b) overviews the neutralizing potency of P4J15 against the different pseudotyped lentiviruses (n=6) and VLPs (n=4).

### Prophylactic use of P4J15 in the hamster Omicron BA.5 infection model

To further validate the potency of P4J15, *in vivo* live virus challenge experiments were performed in a prophylactic hamster challenge model of SARS-CoV-2 infection. Animals were dosed with 5, 1 or 0.5 mg/kg of P4J15, 5 mg/kg of bebtelovimab or a human IgG1 control, challenged two days later with an intranasal inoculation of the Omicron BA.5 SARS-CoV-2 virus (**Fig. 4a**) and lung tissue from the hamsters were examined four days later for infectious virus and viral RNA. Infectious virus was undetectable in the lungs of all but one P4J15 treated hamsters in the lowest dose 0.5 mg/kg group, which still had a >2-log reduction in levels of infectious virus compared to the isotype mAb-treated control animals (**Fig. 4b**). In comparison, 1 out of 5 hamsters in the 5 mg/kg bebtelovimab group had detectable levels of infectious virus. Importantly, protective plasma levels of P4J15 in Omicron BA.5 challenged hamsters were shown to be ∼7 µg/ml, whereas in the bebtelovimab arm of the study, the one treated animal with detectable infection virus in the lungs had mAb plasma levels of 83 µg/ml. Interestingly, although all P4J15 treatment groups showed a significant reduction in genomic RNA levels, relatively high levels were detected in two and three hamsters for the 1 and 0.5 mg/kg dose arms (**Fig. 4c**). This indicates that although P4J15 treatments virtually eliminated infectious virus, RNA, presumably from inactivated virus, was still detectable in select animals four days after challenge.

**Figure 4:**
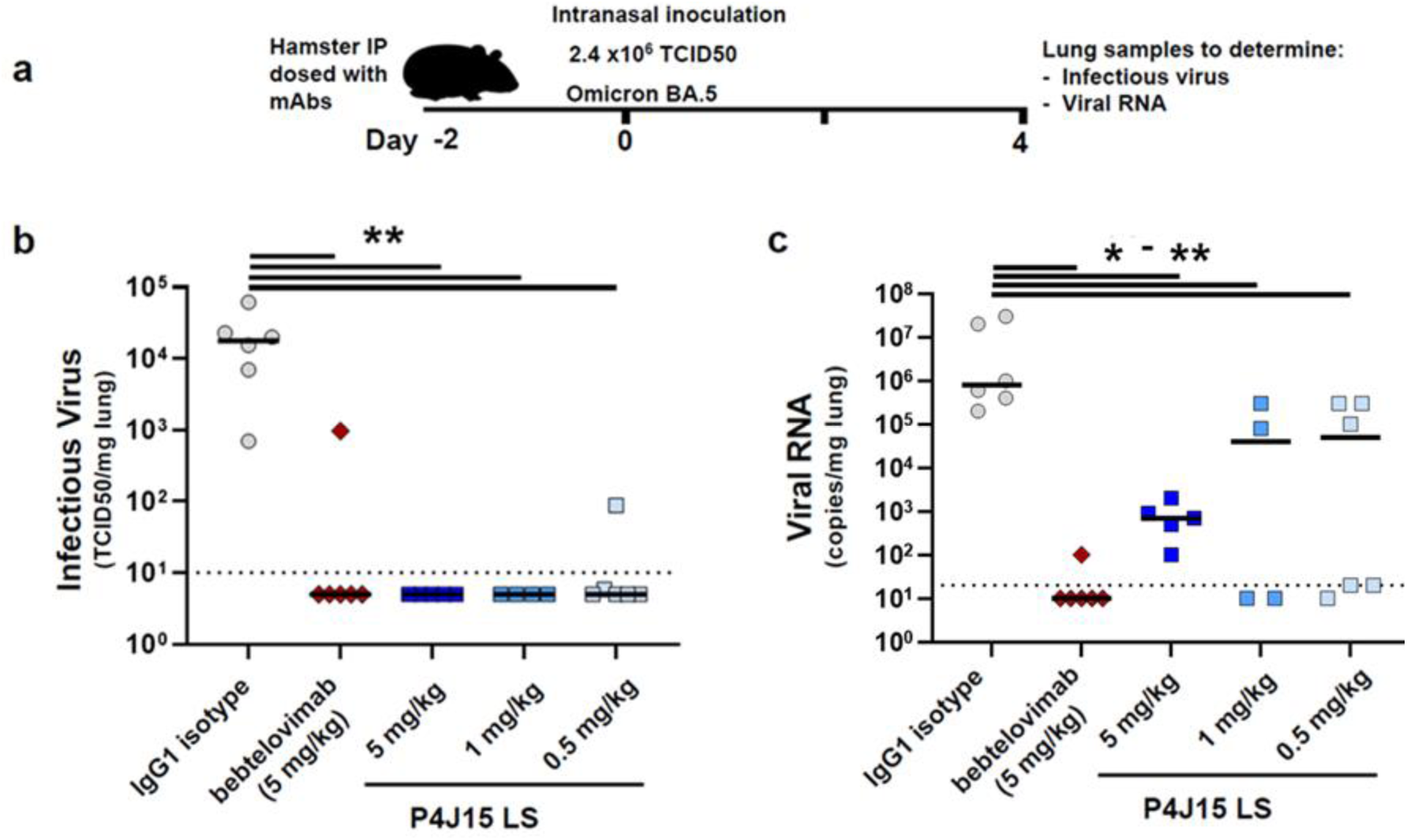
*In vivo* efficacy in the SARS-CoV-2 Omicron BA.5 hamster challenge model. **a**) Overview of study design for the SARS-CoV-2 hamster challenge model. Animals were administered intraperitoneally 5.0, 1.0 or 0.5 mg/kg of P4J15, 5 mg/kg of bebtelovimab positive control or 5 mg/kg of an IgG1 isotype control and challenged two days later (Day 0) with an intranasal inoculation of the Omicron BA.5 SARS-CoV-2 virus (2.4 x10^6^ TCID50). **b**) Median levels of infectious virus and **c**) viral RNA copies/mg lung tissue in each of the study arms are shown for day 4 post-inoculation with SARS-CoV-2 virus. A total of 4-6 hamsters were used per P4J15 treatment arm. Non-parametric Mann-Whitney two-tailed tests were used to evaluate the statistical difference between the treatment conditions with P= 0.0043, 0.0043, 0.0095 and 0.0022 (**) in b (left to right) and P=0.0022, 0.0043, 0.0190 and 0.0087 (* to **) in c (left to right).

### P4J15 shows full prophylactic therapeutic efficacy in cynomolgus macaques

We next evaluated P4J15 LS with M428L/N434S half-life extension mutations in the Fc domain in mediating protection from live SARS-CoV-2 Omicron XBB.1.5 virus infection in a pre-exposure challenge study in cynomolgus macaques. Non-human primates (NHP, n=6) were administered 5 mg/kg of P4J15 intravenously and challenged 72 hours later with 1×10^5^ TCID50 of SARS-CoV-2 Omicron XBB.1.5 virus via a combined intranasal and intratracheal route (**Fig. 5a**). Following viral challenge, control animals (n=4, tested in parallel and n=2 historical controls) showed similar genomic (g)RNA levels and kinetics with median peak viral loads (VL) of 7.3, 7.1 and 6.4-log10 copies/ml gRNA in nasopharyngeal swabs, tracheal swabs and bronchoalveolar lavage (BAL) samples, respectively, at 2-3 days post challenge (**Fig. 5b-d**). In comparison, the six P4J15 LS treated NHPs had essentially undetectable levels of gRNA at all sample and time points tested. This complete protection provided by P4J15 LS was further confirmed by evaluating signs of active viral replication, as assessed by subgenomic (sg)RNA levels, which peaked in control animals at 3-4 days post-challenge with nasopharyngeal swabs, tracheal swabs and BAL showing median values of 5.2- 5.2 and 4.2-log10 copies per ml, respectively (**Fig. 5e-g**). As expected with the almost complete viral suppression, area under the curve analysis (AUC) for P4J15 LS treated NHPs showed a strongly significant reduction in gRNA compared to controls in nasopharyngeal samples collected from left and right nostrils throughout the study and tracheal samples (P<0.0001 and 0.0022, respectively) (**Fig. 5h**). Similar strong and significantly reduced levels of sgRNA in the AUC analysis was observed in P4J15 LS treated NHPs where sgRNA was undetected in all samples analyzed throughout the study period (**Fig. 5i**). This indicates the absence of any replicating virus and the complete prophylactic protection provided by P4J15 LS in a non-human primate model.

**Figure 5:**
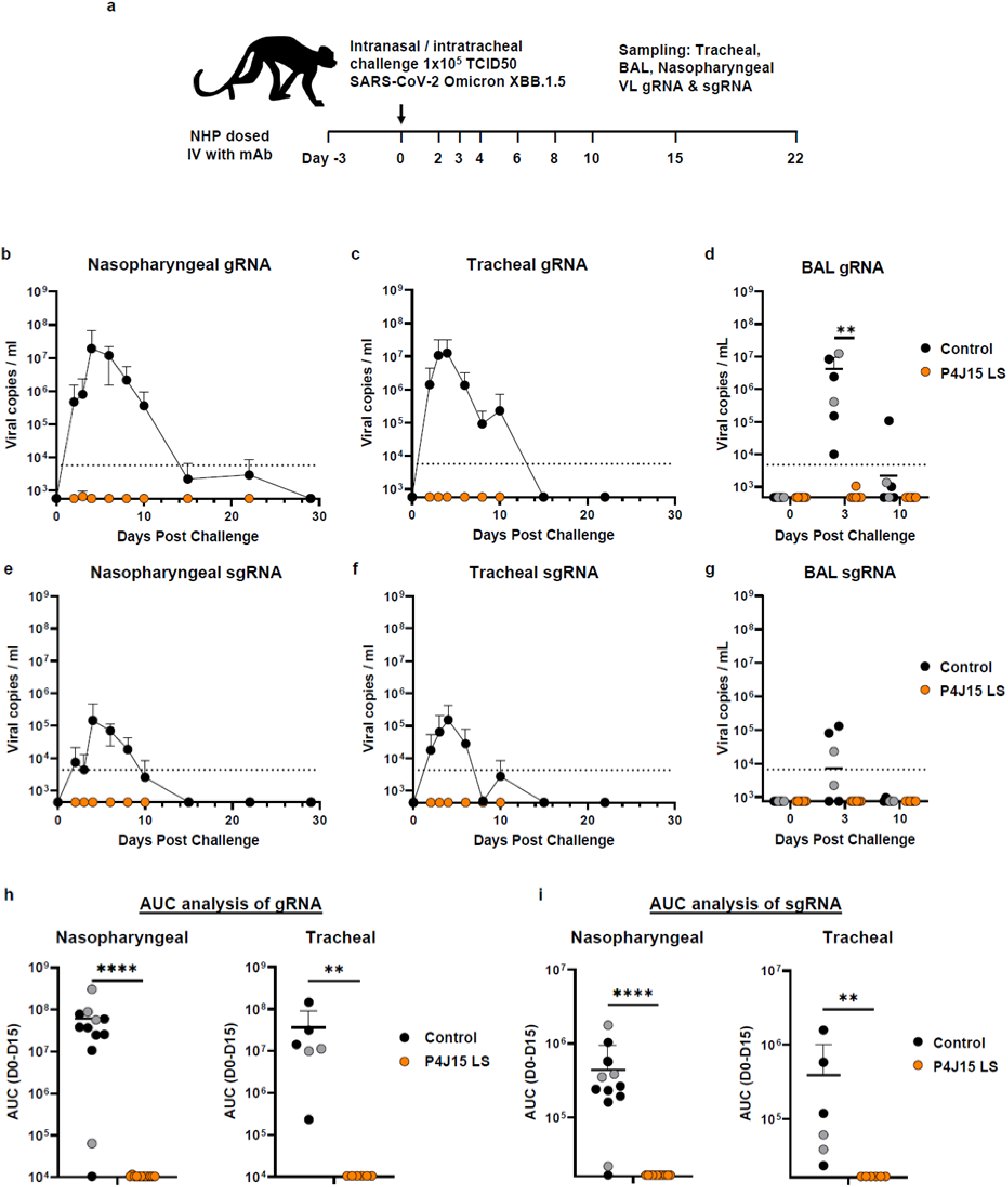
*in vivo* efficacy of P4J15 against the SARS-COV-2 XBB.1.5 virus infection in the non-human primate (NHP) challenge model. **a)** Overview of study design for the SARS-CoV-2 NHP challenge model. Six animals were administered intravenous 5 mg/kg of P4J15 LS and challenged three days later (Day 0) along with four control animals (in black) via intranasal and intratracheal inoculation of the Omicron XBB.1.5 SARS-CoV-2 virus (1×10^5^ TCID50). Tracheal fluids, nasopharyngeal fluids and bronchoalveolar lavages (BAL) collected during the course of the study were evaluated for viral copies per ml of genomic (g)RNA **b)-d)** and subgenomic (sg)RNA **e)-g** with data plotted to include two historical control animals (grey circles) infected with the same inoculum and batch of Omicron XBB.1.5 virus. **h)** Area under the curve (AUC) analysis of gRNA detected between days 0 and 15 of the study. Individual data for nasopharyngeal fluids collected from left and right nostril for each of the timepoints (n=12) and tracheal fluids (n=6) were plotted in left and right panels, respectively. **i**) AUC analysis of sgRNA detected between days 0 and 15 of the study for samples as indicated in h). Mean values ± SD are shown, and Mann-Whitney two-sided tests were performed to compare study groups in panels d, h, i with P values of 0.0022 (**) for d; and p<0.0001 (****) and 0.0022 (**) of h and i. Lower limit of detection were 2.76- and 2.63-log copies per ml for viral gRNA and sgRNA, respectively. Dotted line indicates lower limit of quantitation at 3.76- and 3.63-log copies per ml for viral gRNA and sgRNA, respectively.

## DISCUSSION

As the WHO declares that the emergency phase of the SARS-CoV-2 pandemic has ended, the strain on the health care system continues with hospitalization rates from new infections still reaching >15’000 patients per week across North America and Europe (Supplementary Table 1). Furthermore, it was recently reported that 10% of individuals suffer from long COVID after Omicron infection, with clinical symptoms that can include fatigue, brain fog, and dizziness lasting for upwards of six months (*21*). Newly emerged SARS-CoV-2 variants, including BQ.1.1 and XBB variants, now up to XBB.2.3, are increasingly infectious and immune evasive, significantly eroding the protection conferred by vaccines and previous infections. In addition, almost all licensed monoclonal antibodies for SARS-CoV-2 are inactive against currently circulating VOCs.

Here we report the isolation of the fully human P4J15 antibody from a vaccinated, post-infected donor with superior breadth and neutralizing potency to all other reported antibodies against SARS-CoV-2 VOCs up to the most recent XBB.2.3 (*8–11*). This unique antibody has been extensively optimized *in vivo* by somatic hypermutation, as evidenced by the low 82.5% and 90.5% identity of the P4J15 heavy and light chain sequences, respectively, relative to the IGHV4-34*01 and IGKV1-33*01 germline genes. The uniqueness of P4J15 is also illustrated by the low identity of 53.3% with the closest match of the 12016 anti-spike HCDR3 sequences reported to date.

Cryo-EM performed with the antibody Fab bound to Omicron XBB.1 Spike revealed that P4J15 binds as a Class 1 mAb to the up-RBD conformation of the Spike trimer with a large, buried surface area of 828 Å^2^. Importantly, of the 26 RBD residues that make up the P4J15 binding epitope, 22 are shared with those used for ACE2 binding, representing 93% of the P4J15 contact site. Conversely, ∼87% of the 887 Å^2^ ACE2 binding epitope on RBD is shared by P4J15, which accounts for its potent neutralizing mechanism of action through blockade of ACE2 binding to all SARS-CoV-2 Spike variants tested to date. Interestingly, there are 10 RBD residues shared between P4J15 and ACE2 that have undergone evolutionary fine-tuning to evade neutralizing antibody responses while optimizing ACE2 binding and/or viral infectivity. Through the pandemic, these substitutions relative to the ancestral Spike include S477N, T478K/R, E484K/A, F486V/S/P, F490S, Q493R/Q, G496S, Q498R, N501Y and Y505H. Selection of these mutations over the last 42 months has contributed to the XBB.1 and BQ.1.1 variants exhibiting a 7.6- and 17-fold increase in infectivity relative to Omicron BA.2, respectively, as monitored in pseudovirus and cell fusion assays (*22, 23*). Indeed, similar to ACE2 and consistent with having ACE2 mimetic properties, we see that P4J15 has improved neutralizing activities against post-Omicron variants (EC50 values of 5 to 14 ng/ml) compared to the ancestral 2019-nCoV (EC50 of 41 ng/ml) in the Spike pseudotyped lentiviral neutralization assays.

Even with the highly overlapping binding epitope of P4J15 and ACE2 on the Spike RBD, we confirmed that the development of resistance is inevitable when a virus is under selective pressure, at least *in vitro*. Mutations centered at F456, A475, G476, N477, N487 and Y489 did confer escape to neutralization by P4J15 but also reduced binding to ACE2 by 1- to 3-logs, indicating that the virus incurs a significant fitness penalty in developing resistance. Although epistasis is always possible to complement for deleterious mutations (*24*), the almost complete absence of these specific resistance mutations within the GISAID database confirms that the virus cannot easily escape the type of inhibition imposed by P4J15 without compromising its ability to spread in the population. Of note, substitutions at some of the incriminated positions are detectable in public sequence databases, albeit at very low frequency, but these mutations do not confer resistance to neutralization by P4J15, which requires very specific amino acid substitutions.

Finally, *in vivo* efficacy studies performed with two animal models demonstrate that P4J15 provides exceptional levels of protection against infection. In hamsters challenged with the Omicron BA.5 virus, animals pre-dosed with 0.5 mg/kg of mAb resulting in serum concentrations as low as 7 µg/ml were strongly protected from infection with almost complete suppression of infectious virus in the lungs. A cynomolgus monkey challenge performed with the latest available XBB.1.5 SARS-CoV-2 variant further revealed that P4J15 at 5 mg/kg produced near sterilizing protection in all six treated animals, with only two positive samples with viral genomic RNA near the detection limit of the RT-PCR assay out of a total of 192 nasal, tracheal and BAL samples tested. These results are among the most definitive protection results for SARS-CoV-2 non-human primate challenge studies (*25–27*). It should also be noted that P4J15 used in these *in vivo* studies was produced with the LS extended half-life mutations in the Fc domain, which can provide upwards of six months prophylactic protection following a single dose of antibody administer by intravenous, intramuscular or intraperitoneal injection (*28, 29*). With its impressive *in vivo* protection, P4J15 LS could form a strong basis for prophylactic therapy in immunocompromised patients. To supplement this activity and protect against the development of resistance (*30*), P4J15 could potentially be combined as a cocktail with a second neutralizing antibody that binds a distinct epitope on Spike. For example, although sotrovimab is only weakly to moderately active against some of the newer SARS-CoV-2 variants, this antibody would help to suppress any low-level, poorly fit viral quasi-species that are resistant to P4J15.

The world is at a critical juncture in the SARS-CoV-2 pandemic where humoral protection afforded by vaccines is waning, the public at large has become complacent and new variants with ever-increasing infectivity and immune resistance are emerging regularly. As a result of these factors, the most vulnerable in our population, the immunocompromised, those with comorbidities such as cancer and the elderly, who are unable to mount a strong protective humoral immune response after vaccination (*31*), are at significantly increased risk of hospitalization and death. Given the potent neutralizing activity of P4J15, its ACE2 mimetic properties that may help to limit the development of resistant virus and the impressive complete *in vivo* protection in the non-human primate model, this monoclonal antibody has the potential to be a superior anti-SARS-CoV-2 mAb for prophylactic and/or potentially therapeutic interventions. Furthermore, the breath of potent neutralizing activity against all current VOCs and variant quasi-species within public SARS-CoV-2 sequence databases suggests that it may be able to neutralize many future VOCs that could emerge in the months and years ahead, providing a sustainable and long-lasting solution to protect the most vulnerable in our population.

## Supplementary Figures

**Supplementary data Fig. 1.**
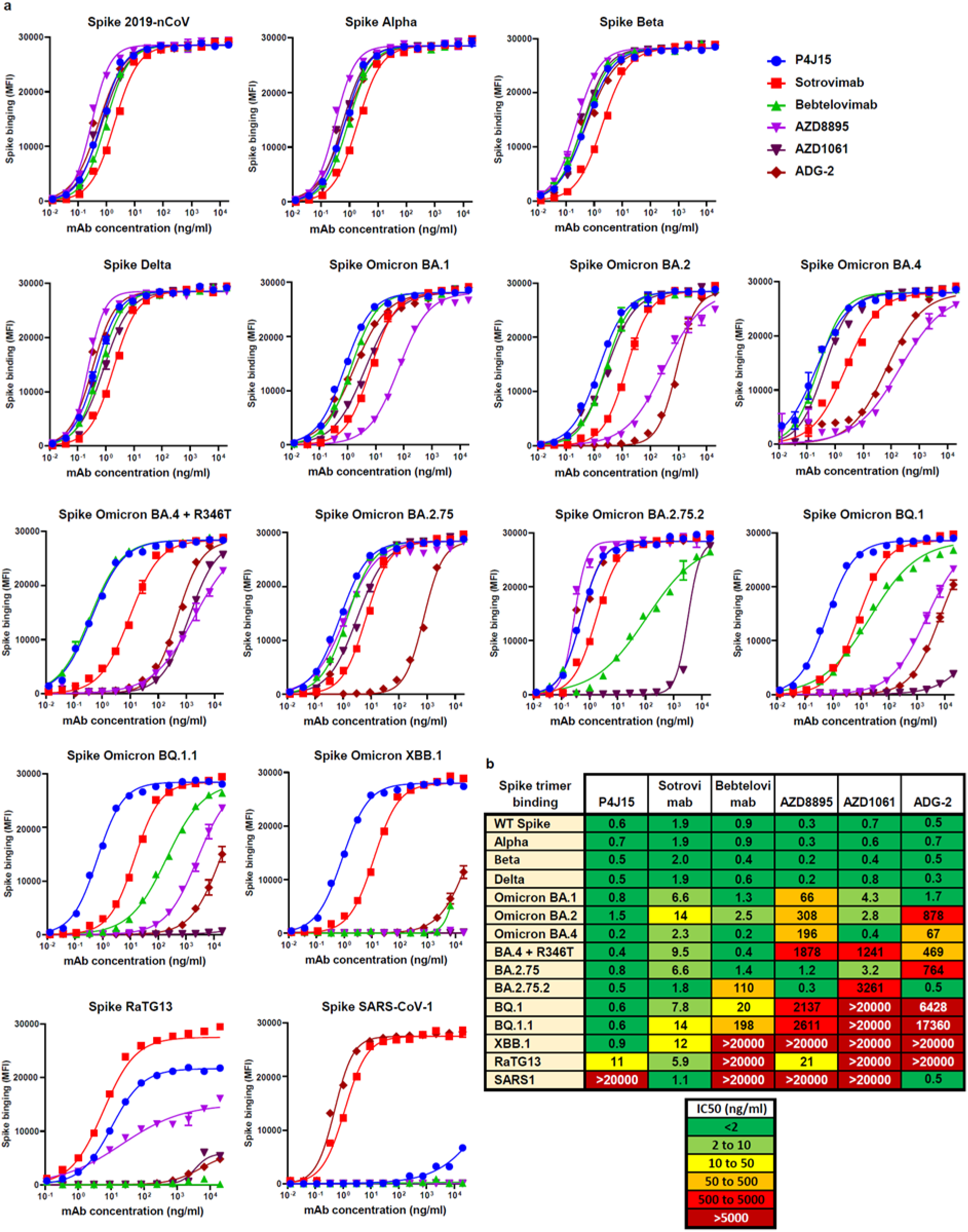
Binding properties of P4J15 and other anti-SARS-CoV-2 antibodies for recombinant Spike trimer proteins from SARS-CoV-2 2019-nCoV to Omicron XBB.1, and sarbecovirus RaTG13 and SARS-CoV-1 proteins. **a)** Spike binding curves performed in a Luminex bead-based assay. **b)** Heatmap table showing binding affinity IC50 values for our panel of mAbs to the indicated Spike trimer proteins. Data presented are representative of 2-4 independent experiments with each concentration response tested in duplicate. Mean values ± SEM are shown.

**Supplementary data Fig. 2.**
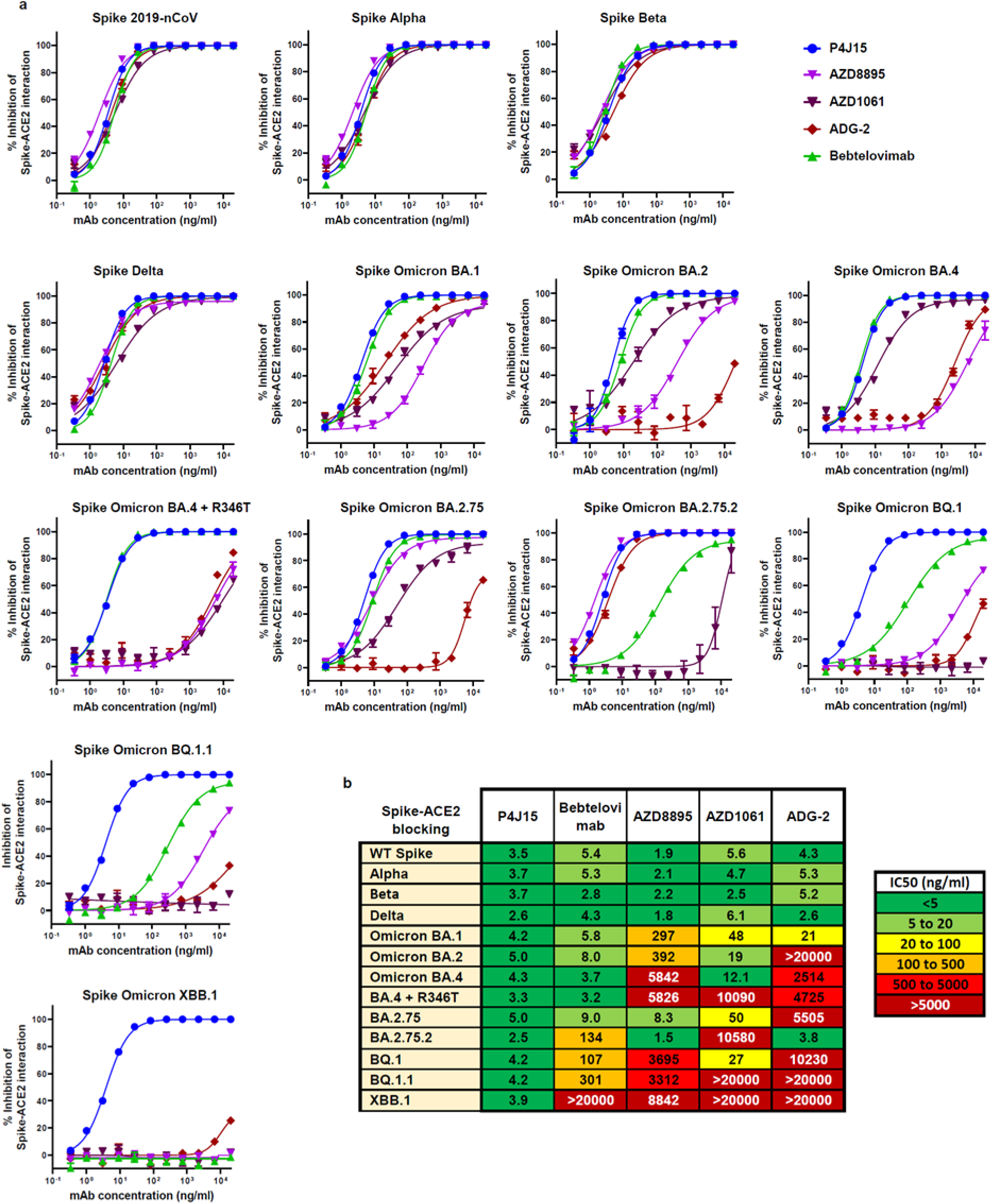
P4J15 is the most potent and broadly active antibody in a Spike-ACE2 surrogate neutralization assay performed with trimeric Spike proteins from a panel of SARS-CoV-2 variants of concern. **a)** Spike-ACE2 blocking activity of P4J15 compared to a panel of authorized and clinically advanced anti-Spike mAbs. **b)** Heatmap table showing IC50 values for our panel of mAbs in the Spike-ACE2 assay. Luminex based assays were performed with beads coupled with Spike trimer proteins from the original 2019-nCoV SARS-CoV-2, Alpha, Beta, Gamma, and the different Omicron lineages listed. Sotrovimab was not included in this analysis as it binds the RBD without blocking the Spike-ACE2 interaction. Data presented is representative of 2-4 independent experiments with each concentration response tested in duplicate. Mean values ± SEM are shown.

**Supplementary data Fig. 3.**
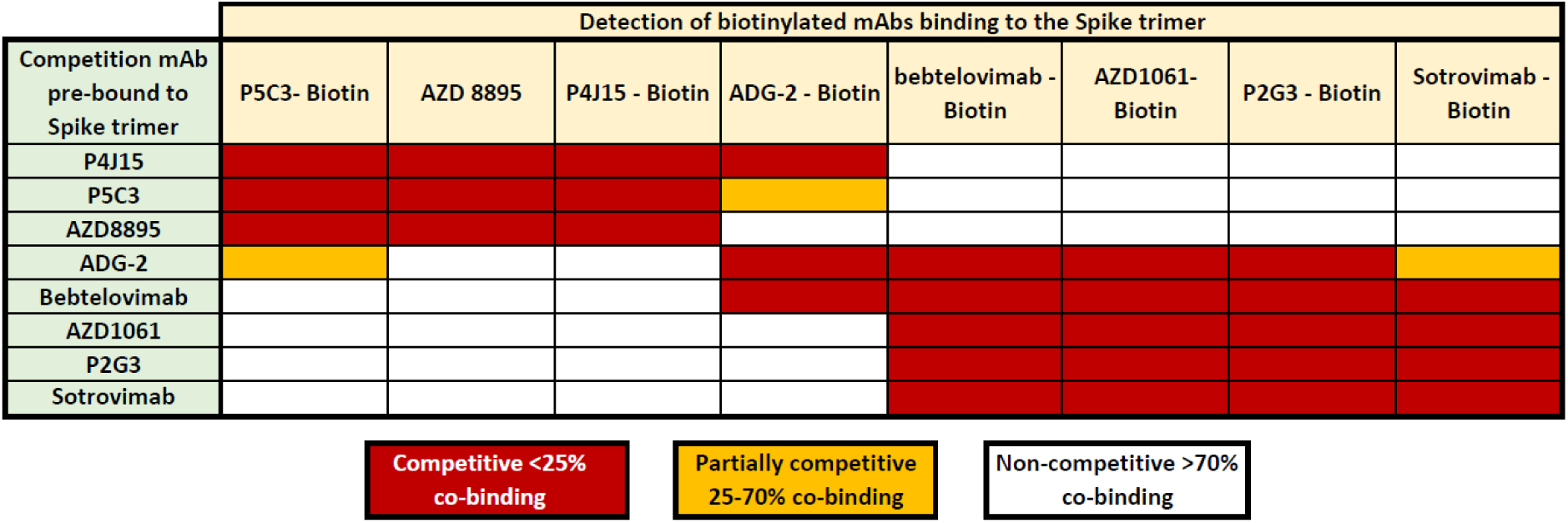
P4J15 binds competitively to the Spike trimer with Class 1 neutralizing antibodies. Competitive binding studies between antibodies binding to the 2019-nCoV Spike trimer protein. Spike coupled beads pre-incubated with saturating concentrations of competitor antibody were used for binding studies with the indicated biotinylated antibodies. Competitors induced either strong blocking (Red boxes), partial competition (orange boxes) or non-competitive (white boxes) binding with the corresponding antibody to Spike.

**Supplementary Figure 4.**
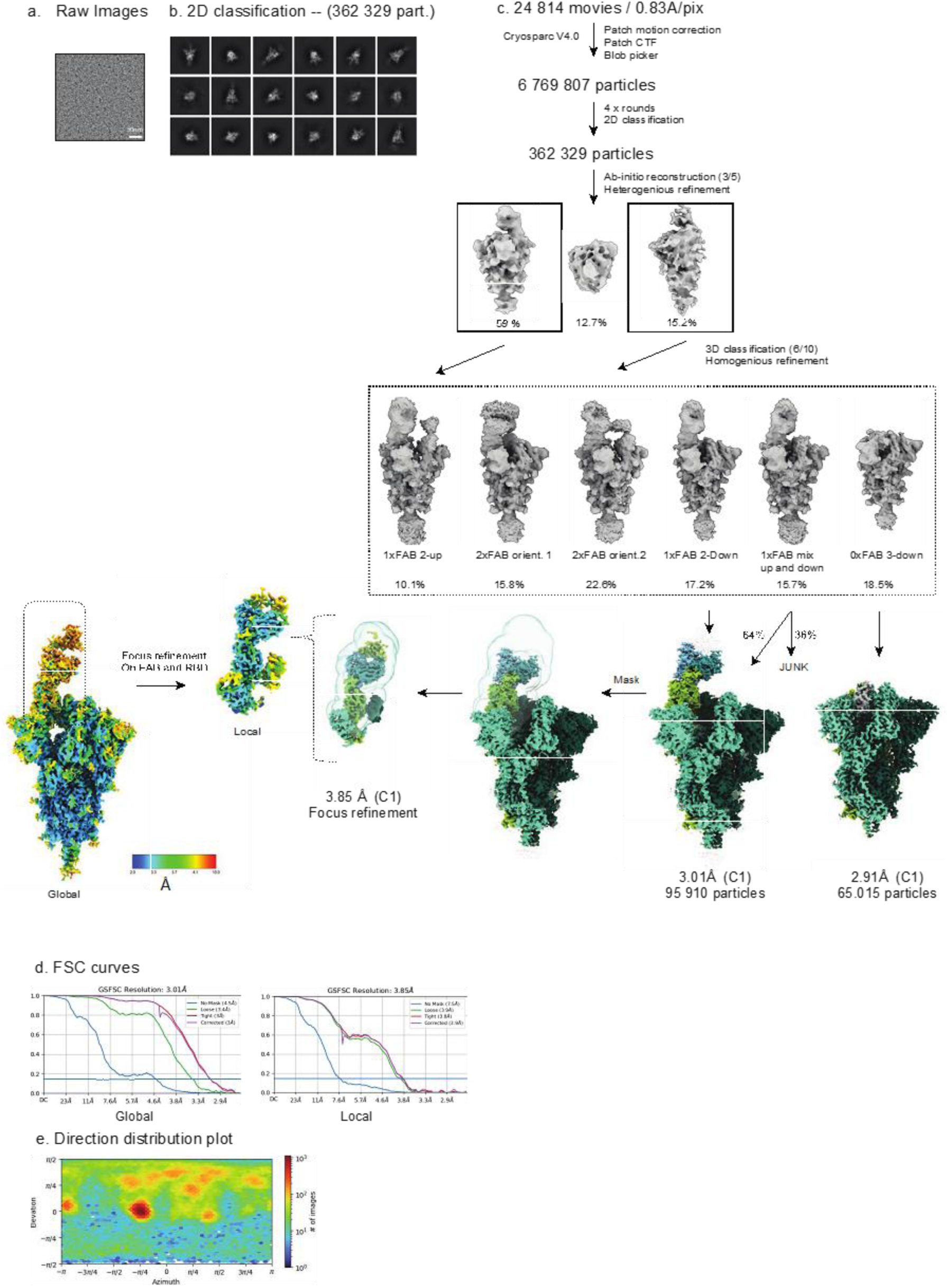
Details of Cryo-EM processing and Resolution maps. **a)** Raw representative micrograph. **b)** Representative 2D class averages. **c)** Cryo-EM processing workflow performed in CryoSPARC **d)** FSC curves indicating a resolution of 3.01 Å of the full-length Omicron XBB.1 Spike bound to the P4J15 Fab and 3.85 Å for the focused local refinement, **e)** Direction distribution plot

**Supplementary Figure 5.**
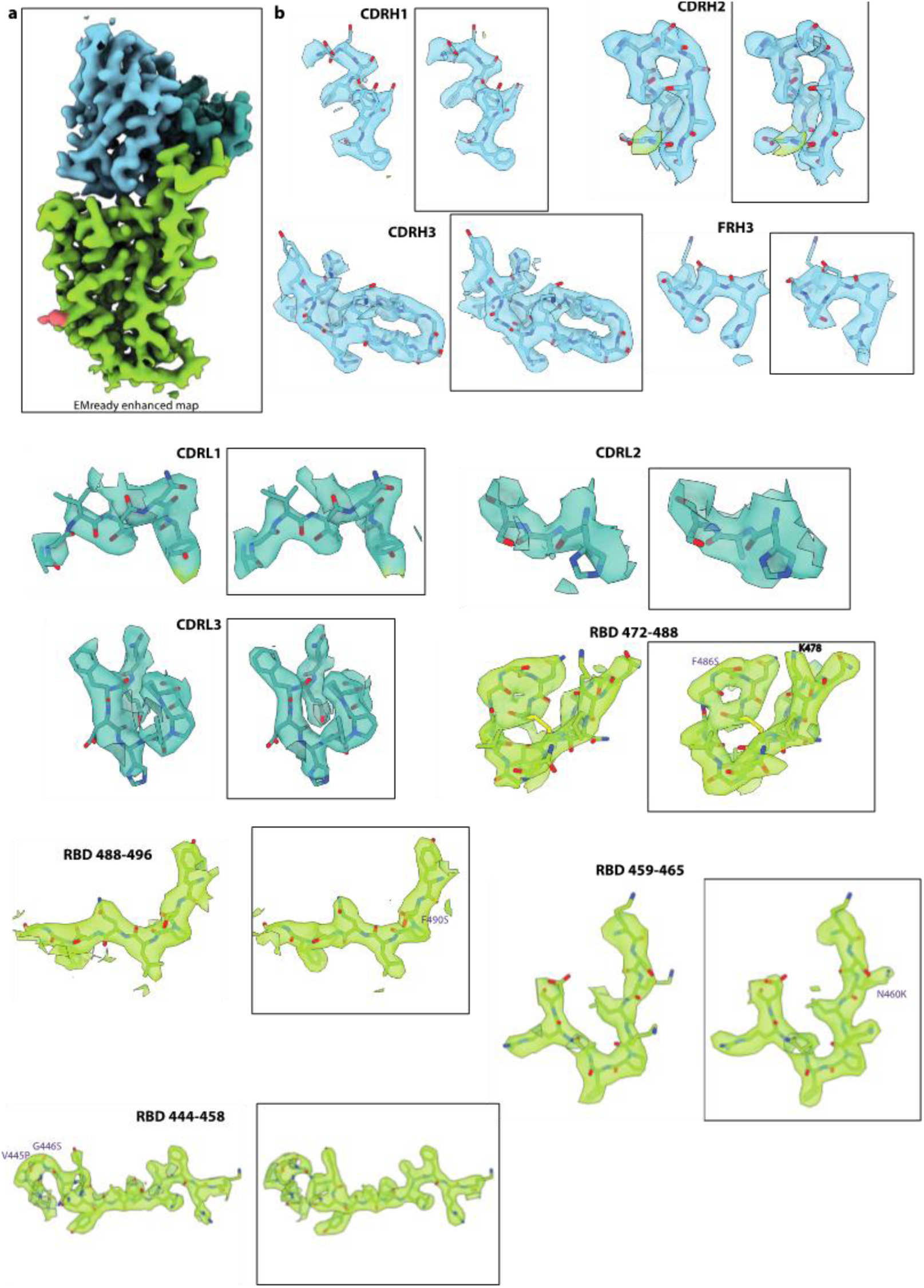
Highlights of regions of the XBB.1 RBD and P4J15 with Cryo-EM density maps from Cryosparc and from EMReady. The Cryo-EM density is rendered as a mesh. The atomic model is shown as ribbon or stick representation. Representations in a box are maps from EMReady.

**Supplementary Figure 6.**
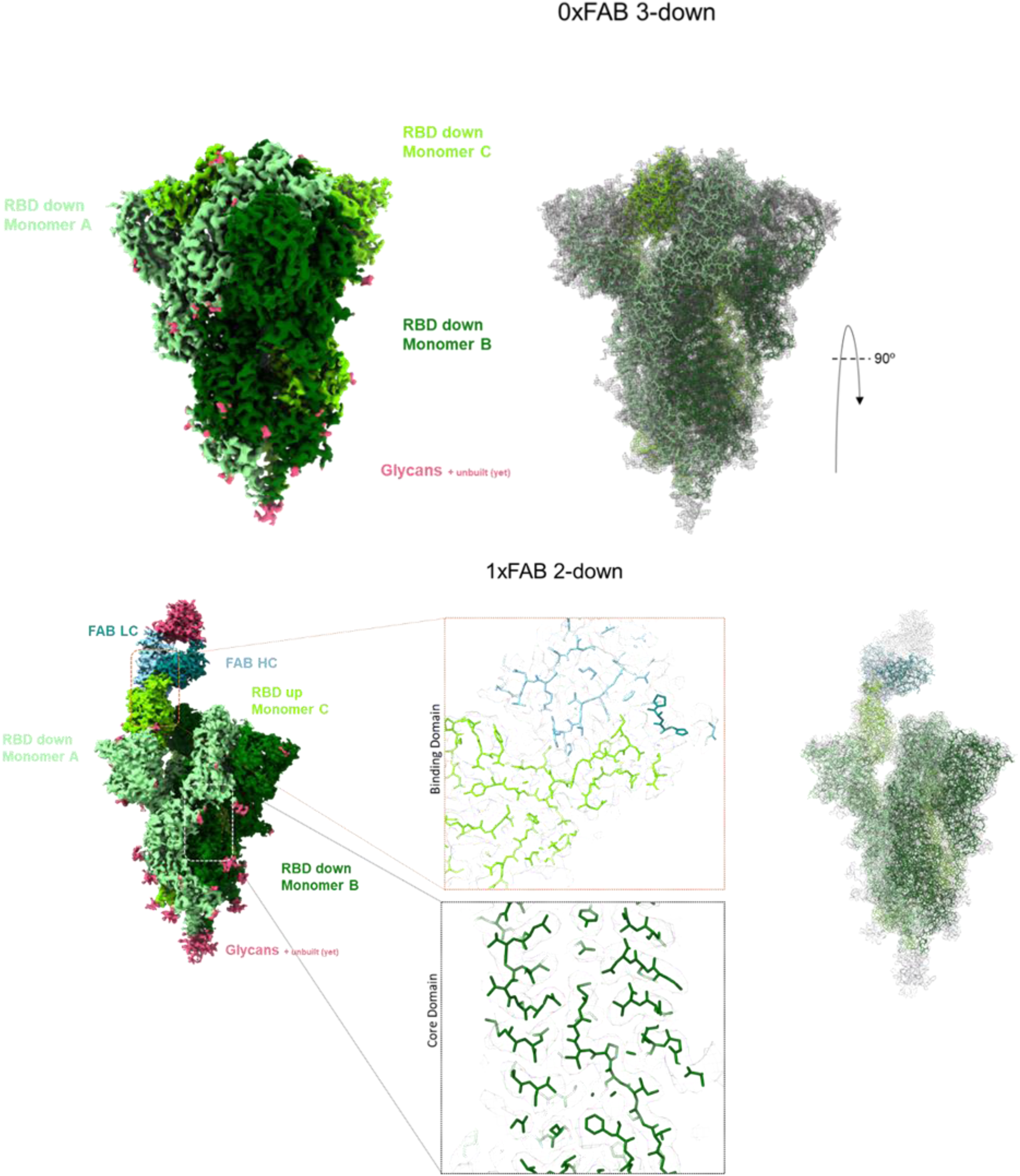
Details of Spike XBB.1 trimer 3D classifications. **a)** Spike trimer with all three RBDs in the down conformation at 2.91 Å resolution and **b)** Spike trimer with P4J15 Fab bound in the RBD up conformation and the remaining two RBD monomers in the down conformation.

**Supplementary Figure 7.**
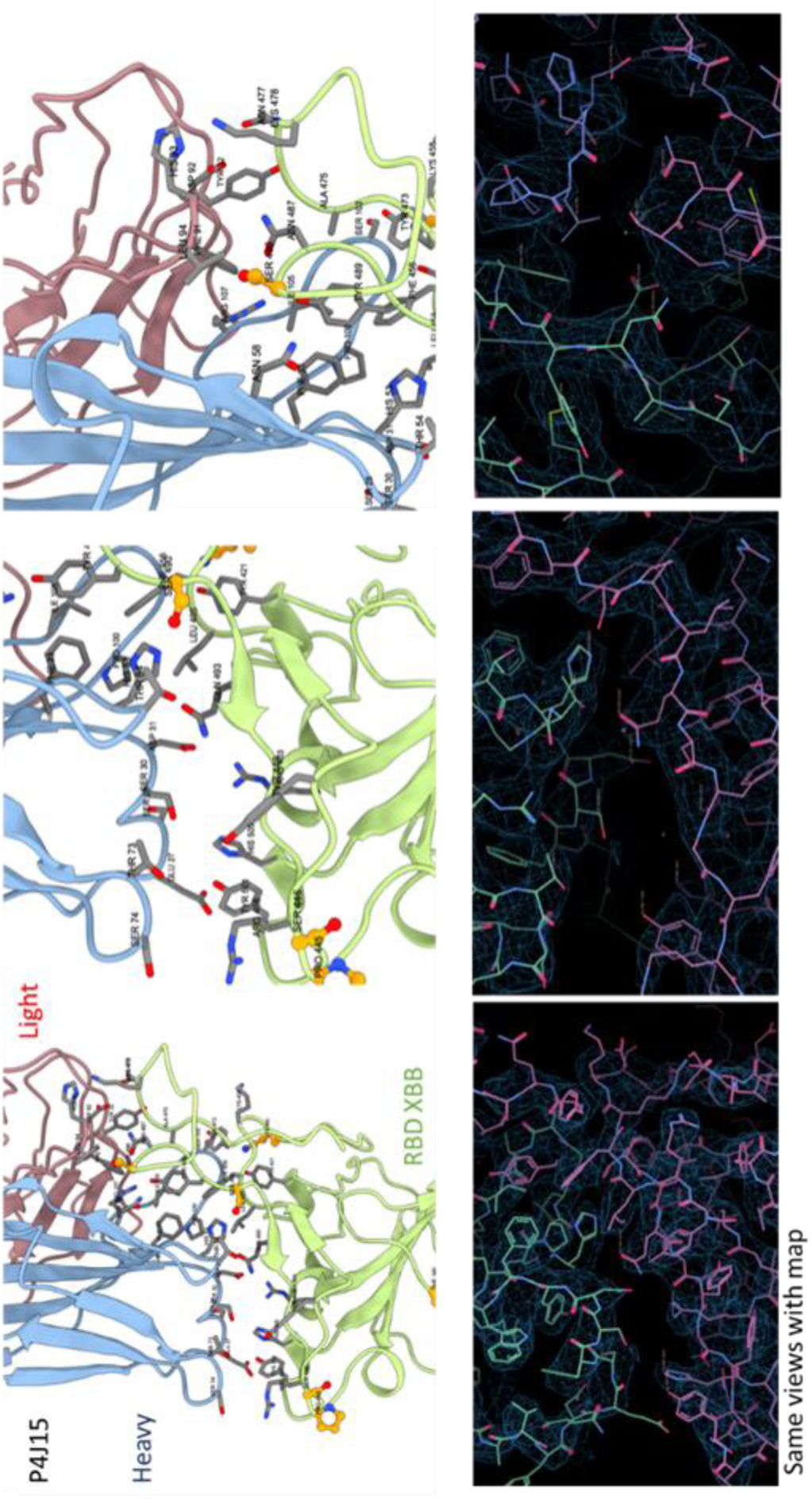
Additional views of the P4J15 Fab-Omicron XBB.1 RBD interactions. Ribbon structure with stick representation of contact residues for P4J15 and RBD from three different views (top panels) with corresponding mesh representation of Cryo-EM density (bottom panels).

**Supplementary Figure 8.**
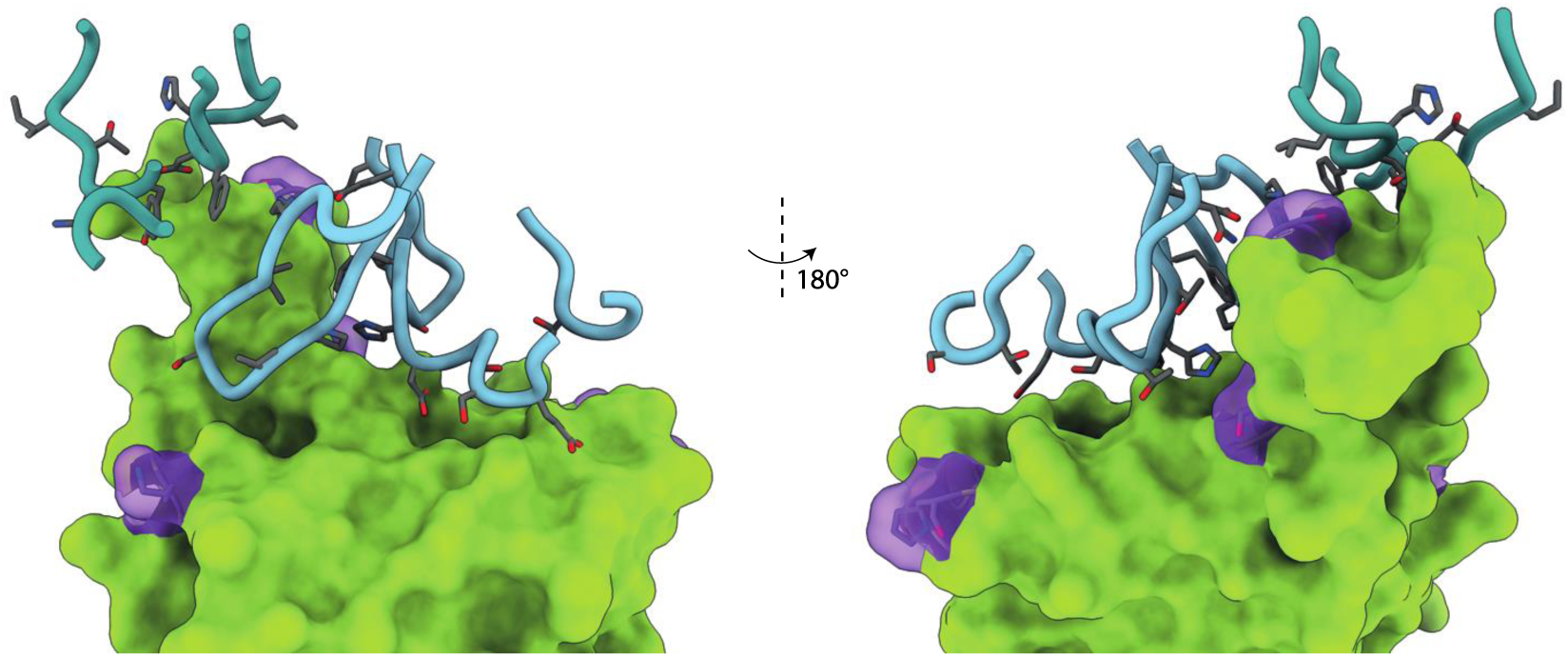
Back and front view representation of the RBD (green) and P4J15 heavy and light chain contact loops shown in cyan and turquois.

Supplementary Table 1: Estimated new hospitalization rates in the United States and Europe.

| Country / Region | Average weekly new hospitalizations: May 1<sup>st</sup> to June 18<sup>th</sup>, 2023, from Our World in Data | Population | New hospitalizations per week per 100000 individuals |
| --- | --- | --- | --- |
| Belgium | 248 | 11'590'000 | 2,14 |
| Czechia | 81 | 10'709'000 | 0,76 |
| Denmark | 210 | 5'792'000 | 3,62 |
| England | 2133 | 55'977'000 | 3,81 |
| Estonia | 41 | 1'327'000 | 3,08 |
| Germany | 1330 | 83'784'000 | 1,59 |
| Greece | 500 | 10'423'000 | 4,80 |
| Hungary | 36 | 9'660'000 | 0,37 |
| Ireland | 207 | 4'938'000 | 4,20 |
| Italy | 777 | 60'462'000 | 1,29 |
| Latvia | 74 | 1'886'000 | 3,92 |
| Northern Ireland | 36 | 1'903'000 | 1,87 |
| Slovakia | 44 | 5'460'000 | 0,81 |
| Spain | 1867 | 46'755'000 | 3,99 |
| Wales | 4 | 3'210'000 | 0,13 |
| Data available for Europe countries | 7587 | 313'875'196 | 2,42 |
| United States | 8028 | 336'862'000 | 2,38 |
| **Weekly new hospitalizations from Northern America and Europe** | **15615** |  |  |

Supplementary Table 2: SARS-CoV-2 Spike variant substitutions.

**Supplementary Table 2: SARS-CoV-2 Spike variant substitutions**
|  |  |
| --- | --- |
| **D614G** | D614G |
| **Alpha**<br>**B.1.1.7** | Δ69-70, Δ144, N501Y, A570D, D614G, P681H, T716I, S982A, D1118H |
| **Beta**<br>**B.1.351** | L18F, D80A, D215G, Δ242-244, R246I, K417N, E484K, N501Y, D614G, A701V |
| **Gamma**<br>**P.1** | L18F, T20N, P26S, D138Y, R190S, K417T, E484K, N501Y, D614G, H655Y, T1027I, V1176F |
| **delta**<br>**B.1.617.2** | T19R, Δ156-157, R158G, L452R, T478K, D614G, P681R, D950N |
| **delta**<br>**AY.4.2** | T19R, T95I, Y145H, Δ156-157, R158G, A222V, L452R, T478K, D614G, P681R, D950N |
| **Kappa**<br>**B.1.617.1** | E154K, L452R, E484Q, D614G, P681R, Q1071H |
| **Lambda**<br>**C.37** | G75V, T76I, R246N, del247-253, L452Q, F490S, D614G, T859N |
| **Iota**<br>**B.1.526** | L5F, T95I, D253G, E484K, D614G, A701V |
| **Eta**<br>**B.1.525** | Q52R, del69-70, E484K, Q677H, F888L |
| **Omicron**<br>**BA.1** | A67V, Δ69-70, T95I, G142D, Δ143-145, Δ211, L212I, ins214EPE, G339D, S371L, S373P, S375F, K417N, N440K, G446S, S477N, T478K, E484A, Q493R, G496S, Q498R, N501Y, Y505H, T547K, D614G, H655Y, N679K, P681H, N764K, D796Y, N856K, Q954H, N969K, L981F |
| **Omicron**<br>**BA.1.1** | A67V, Δ69-70, T95I, G142D, Δ143-145, Δ211, L212I, ins214EPE, G339D, R346K, S371L, S373P, S375F, K417N, N440K, G446S, S477N, T478K, E484A, Q493R, G496S, Q498R, N501Y, Y505H, T547K, D614G, H655Y, N679K, P681H, N764K, D796Y, N856K, Q954H, N969K, L981F |
| **Omicron**<br>**BA.2** | T19I, Δ24-26, A27S, G142D, V213G, G339D, S371F, S373P, S375F, T376A, D405N, R408S, K417N, N440K, S477N, T478K, E484A, Q493R, Q498R, N501Y, Y505H, D614G, H655Y, N679K, P681H, N764K, D796Y, Q954H, N969K |
| **Omicron**<br>**BA.2.12** | T19I, Δ24-26, A27S, G142D, V213G, G339D, S371F, S373P, S375F, T376A, D405N, R408S, K417N, N440K, S477N, T478K, E484A, Q493R, Q498R, N501Y, Y505H, D614G, H655Y, N679K, P681H, S704L, N764K, D796Y, Q954H, N969K |
| **Omicron**<br>**BA.2.12.1** | T19I, Δ24-26, A27S, G142D, V213G, G339D, S371F, S373P, S375F, T376A, D405N, R408S, K417N, N440K, L452Q, S477N, T478K, E484A, Q493R, Q498R, N501Y, Y505H, D614G, H655Y, N679K, P681H, S704L, N764K, D796Y, Q954H, N969K |
| **Omicron**<br>**BA.2.75** | T19I, Δ24-26, A27S, G142D, K417E, W152R, F157L, I210V, V213G, G257S, G339D, S371F, S373P, S375F, T376A, D405N, R408S, K417N, N440K, G446S, N460K, S477N, T478K, E484A, Q498R, N501Y, Y505H, D614G, H655Y, N679K, P681H, N764K, D796Y, Q954H, N969K |
| **Omicron**<br>**BA.2.75.7** | T19I, Δ24-26, A27S, G142D, K417E, W152R, F157L, I210V, V213G, G257S, G339D, S371F, S373P, S375F, T376A, D405N, R408S, K417N, N440K, G446S, N460K, S477N, T478K, E484A, F486S, Q498R, N501Y, Y505H, D614G, H655Y, N679K, P681H, N764K, D796Y, Q954H, N969K |
| **Omicron**<br>**BA.2.75.2** | T19I, Δ24-26, A27S, G142D, K417E, W152R, F157L, G181V, I210V, V213G, G257S, G339D, R346T, S371F, S373P, S375F, T376A, D405N, R408S, K417N, N440K, G446S, N460K, S477N, T478K, E484A, F486S, Q498R, N501Y, Y505H, D614G, H655Y, N679K, P681H, N764K, D796Y, Q954H, N969K, D1199N |
| **Omicron**<br>**BA.4 and BA.5** | T19I, Δ24-26, A27S, Δ69-70, G142D, V213G, G339D, S371F, S373P, S375F, T376A, D405N, R408S, K417N, N440K, L452R, S477N, T478K, E484A, F486V, Q498R, N501Y, Y505H, D614G, H655Y, N679K, P681H, N764K, D796Y, Q954H, N969K |
| **Omicron**<br>**BF.7** | T19I, Δ24-26, A27S, Δ69-70, G142D, V213G, G339D, R346T, S371F, S373P, S375F, T376A, D405N, R408S, K417N, N440K, L452R, S477N, T478K, E484A, F486V, Q498R, N501Y, Y505H, D614G, H655Y, N679K, P681H, N764K, D796Y, Q954H, N969K |
| **Omicron**<br>**BQ.1** | T19I, Δ24-26, A27S, Δ69-70, G142D, V213G, G339D, S371F, S373P, S375F, T376A, D405N, R408S, K417N, N440K, K444T, L452R, N460K, S477N, T478K, E484A, F486V, Q498R, N501Y, Y505H, D614G, H655Y, N679K, P681H, N764K, D796Y, Q954H, N969K |
| **Omicron**<br>**BQ.1.1** | T19I, Δ24-26, A27S, Δ69-70, G142D, V213G, G339D, R346T, S371F, S373P, S375F, T376A, D405N, R408S, K417N, N440K, K444T, L452R, N460K, S477N, T478K, E484A, F486V, Q498R, N501Y, Y505H, D614G, H655Y, N679K, P681H, N764K, D796Y, Q954H, N969K |
| **Omicron**<br>**XBB.1/XBB.1.9** | T19I, Δ24-26, A27S, V83A, G142D, ΔY144, H146Q, Q183E, V213E, G252V, G339H, R346T, L368I, S371F, S373P, S375F, T376A, D405N, R408S, K417N, N440K, V445P, G446S, N460K, S477N, T478K, E484A, F486S, F490S, Q498R, N501Y, Y505H, D614G, H655Y, N679K, P681H, N764K, D796Y, Q954H, N969K |
| **Omicron**<br>**XBB.1.5/**<br>**XBB.1.9.1** | T19I, Δ24-26, A27S, V83A, G142D, ΔY144, H146Q, Q183E, V213E, G252V, G339H, R346T, L368I, S371F, S373P, S375F, T376A, D405N, R408S, K417N, N440K, V445P, G446S, N460K, S477N, T478K, E484A, F486P, F490S, Q498R, N501Y, Y505H, D614G, H655Y, N679K, P681H, N764K, D796Y, Q954H, N969K |
| **Omicron**<br>**XBB.1.16** | T19I, Δ24-26, A27S, V83A, G142D, ΔY144, H146Q, E180V, Q183E, V213E, G252V, G339H, R346T, L368I, S371F, S373P, S375F, T376A, D405N, R408S, K417N, N440K, V445P, G446S, N460K, S477N, T478R, E484A, F486P, F490S, Q498R, N501Y, Y505H, D614G, H655Y, N679K, P681H, N764K, D796Y, Q954H, N969K |
| **Omicron**<br>**XBB.1.16.1** | T19I, Δ24-26, A27S, V83A, G142D, ΔY144, H146Q, E180V, Q183E, V213E, G252V, G339H, R346T, L368I, S371F, S373P, S375F, T376A, D405N, R408S, K417N, N440K, V445P, G446S, N460K, S477N, T478R, E484A, F486P, F490S, Q498R, N501Y, Y505H, T547I, D614G, H655Y, N679K, P681H, N764K, D796Y, Q954H, N969K |
| **Omicron**<br>**XBB.2.3** | T19I, Δ24-26, A27S, V83A, G142D, ΔY144, H146Q, E180V, Q183E, V213E, D253G, G339H, R346T, L368I, S371F, S373P, S375F, T376A, D405N, R408S, K417N, N440K, V445P, G446S, N460K, S477N, T478R, E484A, F486P, F490S, Q498R, N501Y, Y505H, P521S, T547I, D614G, H655Y, N679K, P681H, N764K, D796Y, Q954H, N969K |
| **Omicron**<br>**EG.1** | T19I, Δ24-26, A27S, V83A, G142D, ΔY144, H146Q, E180V, Q183E, V213E, G252V, G339H, R346T, L368I, S371F, S373P, S375F, T376A, D405N, R408S, K417N, N440K, V445P, G446S, N460K, S477N, T478R, E484A, F486P, F490S, Q498R, N501Y, Y505H, T547I, Q613H, D614G, H655Y, N679K, P681H, N764K, D796Y, Q954H, N969K |

Supplementary Table 3: Cryo-EM data collection and refinement statistics of SPIKE-FAB complex and SPIKE alone.

**Supplementary Table 3: Cryo-EM data collection and refinement statistics of SPIKE-FAB** 657 **complex and SPIKE alone**
|  | **XBB - P4J15**<br>**(Closed)** | **XBB - P4J15**<br>**(Focus)** | **XBB - P4J15 FULL**<br>**MAP** |
| --- | --- | --- | --- |
| **Data collection processing** | Awaiting AUTH | Awaiting AUTH | Awaiting AUTH |
| EMDB ID | 17850 | 17819 | Awaiting AUTH |
| PDB ID | 8PSD | 8PQ2 | - |
| Detector | Falcon IV | Falcon IV | Falcon IV |
| Magnification | 96 000 | 96 000 | 96 000 |
| Voltage (kV) | 300 | 300 | 300 |
| Electron exposure (e-/A2) | 50 | 50 | 50 |
| Defocus range (um) | -0,25 | -0,25 | -0,25 |
| Pixel size (A) | 0,83 | 0,83 | 0,83 |
| Symmetry imposed | C1 | C1 | C1 |
| Initial particle images (no.) | 6 769 807 | 6 769 807 | 6 769 807 |
| Final particle images (no.) | 65 015 | 95 910 | 95 910 |
| Map resolution (A) | 2,91 | 3,85 | 3,01 |
| FSC threshold | 0,143 | 0,143 | 0,143 |
| Extraction box size (A) | 457 | 457 | 457 |
| Fourrier crop to box size (A) | 299 | 299 | 299 |
| Map pixel size (A) | 1,268 | 1,268 | 1,268 |
| **Refinement** |  |  |  |
| Initial models used (PDB codes) | 7q07 | 7q07 | - |
| Model resolution (A) | 2,9 | 3,7 | - |
| FSC threshold | 0,143 | 0,143 | - |
| Map sharpening B factors (A2) | 51,4 | 70,7 | - |
| **Model composition** |  |  |  |
| Non-hydrogen atoms | 48535 | 5969 | - |
| Protein residues | 3076 | 390 | - |
| Nucleotides | 0 | 0 | - |
| Ligands | 36 | 1 | - |
| **B factors (A2)** |  |  |  |
| Protein * | 74,67 | 69,96 | - |
| DNA * | - | - | - |
| **R.m.s. deviation** |  |  |  |
| Bond lenghts (A) | 0,003 | 0,002 | - |
| Bond angles (A) | 0,47 | 0,501 | - |
| Validation |  |  |  |
| MolProbity score | 1,44 | 2,13 | - |
| Clashscore | 2,2 | 5,53 | - |
| Poor rotamers (%) | 1,6 | 2,73 | - |
| *Ramachandran plot* |  |  |  |
| Favored (%) | 95,76 | 91,67 | - |
| Allowed (%) | 4,24 | 8,33 | - |
| Disallowed (%) | 0 | 0 | - |
| *Model-to-data fit* |  |  |  |
| Ccmask | 0,81 | 0,79 | - |
| Ccbox | 0,64 | 0,6 | - |
| Ccpeaks | 0,6 | 0,44 | - |
| CCvolume | 0,79 | 0,66 | - |

## ONLINE METHODS

### Study COVID-19 donors

Serum and blood mononuclear cell samples were from donors participating in the ImmunoCov and ImmunoVax studies performed by the Immunology and Allergy Service, Lausanne University Hospital with all participants being adults of varying ages and having signed informed consent forms for the use of biological samples. Study design and use of subject samples were approved by the Institutional Review Board of the Lausanne University Hospital and the ‘Commission d’éthique du Canton de Vaud’ (CER-VD with trial reference numbers 2020-00620 and 2021-00041, respectively).

### Production of SARS-CoV-2 Spike proteins

SARS-CoV-2 Spike mutations are listed in Supplementary Table 1. Production of 2019-nCoV (D614G), Alpha, Beta, Delta and Omicron BA.1 variants has already been described (*17*). BA.2 and further Omicron sublineages ORFs were cloned by 1kb gBlocks assembly (IDT DNA) and In-Fusion cloning into the nCoV-BA.1 plasmid described earlier (*32*). Single mutations were further introduced by PCR-mediated mutagenesis in each sublineage. The full Omicron ORFs were sequence verified for all the clones. The final constructs encode the Spike ectodomains, containing a native signal peptide, the 2P and furin cleavage site mutations, a C-terminal T4 foldon fusion domain to stabilize the trimer complex followed by C-terminal 8x His and 2x Strep tags for affinity purification. The trimeric Spike variants were produced and purified as previously described (*12*). The purity of Omicron Spike trimers used for cryo-EM was determined to be >98% pure by SDS-PAGE analysis. Biotinylation of Spike or RBD proteins was performed using the EZ-Link™ NHS-PEG4-Biotin (Life Technologies, USA) using a 3- fold molar excess of reagent and using the manufacturer’s protocol. Biotinylated proteins were buffer exchanged with PBS using an Amicon Ultra-0.5 with a 3 kDa molecular weight cut-off. Spike and RBD tetramers were prepared fresh before use and formed by combining biotinylated proteins with PE-conjugated Streptavidin (BD Biosciences, USA) at a molar ratio of 4:1.

### Binding and ACE2 blocking studies with SARS-CoV-2 Spike

Luminex beads used for the serological and purified antibody binding assays were prepared by covalent coupling of SARS-CoV-2 proteins with MagPlex beads using the manufacturer’s protocol with a Bio-Plex Amine Coupling Kit (Bio-Rad, France). Each of the SARS-CoV-2 Spike proteins expressed with different mutations were coupled with different colored MagPlex beads so that tests could be performed with a single protein bead per well or in a multiplexed Luminex binding assay. Binding curves for antibody affinity measurements and the Spike-ACE2 interaction assay were performed as previously described (*12, 33*) using anti-human IgG-PE secondary antibody (OneLambda ThermoFisher; Cat # H10104; 1:100 dilution) for antibody detection in Spike Luminex binding assay and anti-mouse IgG-PE secondary antibody (OneLambda ThermoFisher; Cat# P-21129; 1:100 dilution) in the Spike-ACE2 surrogate neutralization assay. Competitive binding studies were performed by pre-incubating 25 µg/ml of the indicated competitor antibody with the original 2019-nCoV Spike trimer protein coupled Luminex beads for 30 minutes. Biotinylated P4J15, P2G3, AZD8895, AZD1061, bebtelovimab, ADG-2 or sotrovimab antibodies (prepared as described above) were added to each well at 1 µg/ml followed by a further 20-minute incubation. Biotinylated antibody bound to RBD in the presence of competitor was stained with Streptavidin-PE at a 1:1000 dilution (BD Biosciences) and analyzed on a 200 Bioplex instruments. COVID-19 serum samples from 20 donors were monitored for levels of IgG antibody binding to the SARS-CoV-2 Spike trimer proteins from 2019-nCoV, D614G, Alpha, Beta, Delta, Omicron BA.1 and BA.4 in the Luminex bead-based assay.

### Anti-Spike B cell sorting, immortalization and cloning

The blood from a ImmunoVax study donors were collected in EDTA tubes and the isolation of blood mononuclear cell was performed using Leucosep centrifuge tubes (Greiner Bio-one) prefilled with density gradient medium (Ficoll-PaqueTM PLUS, GE Healthcare) according to the manufacturer’s instructions. Freshly isolated cells were stained with the cocktail of fluorescent conjugated antibodies containing mouse anti-human CD19 APC-Cy7 (BD Biosciences; Cat#557791; Clone SJ25C1; 5 µl titration), mouse anti-human CD3-BV510 (BD Biosciences; Cat#563109; Clone UCHT1; 1 µl titration), mouse anti-human IgM-FITC (Biolegend; Cat#314506; clone MHM-88; 2 µl titration), mouse anti-human IgD PE-CF594 (BD Biosciences; Cat#562540; Clone IA6-2; 3 µl titration), mouse anti-human CD27-APC (BD Biosciences; Cat#558664; Clone: M-T271; 5 µl titration), mouse anti-human CD38-V450 (BD Biosciences; Cat#646851; Clone HB7; 5 µl titration) mAbs were used for antigen specific B cell sorting along with the pre-complexed Omicron BA.1 variant Spike tetramer (2 µg in 100µl) coupled to PE-streptavidin (BD Biosciences; Cat#SA10044; 4:1 molar ratio). All other aspects with cell sorting, immortalization protocol using EBV positive supernatants form B95-8 cells and cloning were as described in Fenwick et al (*17*). Sequences for mAbs P4J15 are provided in PDB submissions PDB-8PQ2 and EMD-17819.

### SARS-CoV-2 live virus stocks

All biosafety level 3 procedures were approved by the Swiss Federal Office of Public Health. The SARS-CoV-2 BA.2.75.2 (EPI_ISL_14795784) and BQ.1 (EPI_ISL_15369810) isolates were a kind gift from I. Eckerle, Geneva University Hospitals. Viral stocks were prepared in EpiSerf (ThermoFisher Scientific, USA) with a single passage on VeroE6 cells, aliquoted, frozen, titrated on VeroE6 cells by conventional plaque forming units and sequence verified. BA.2.75.2 isolate differed from our cloned BA.2.75.2 ORF by 1 supplemental mutation (G181V) already found in the original virus isolated from the patient.

### Selection of resistant virus in presence of mAbs

The day before infection, HEK293T ACE2/TMPRSS2 cells previously described (*18*) were seeded in 6-well plates coated with poly-L-lysine at a density of 1.x10^6^ cells per well. To generate a viral population under mAb pressure, early passage virus was diluted in 1.5 ml EpiSerf 2% FCS and incubated with 0.5 ng/ml mAb for 1 hr at 37°C in duplicates. Each mixture was added to the cells and P1 (passage 1) supernatants were harvested 3 days later, clarified on 0.45 µm SPIN-X centrifuge tube filters at 4000×g for 4 minutes. Aliquots of cleared P1 supernatants were diluted 1:40 in DMEM 2%, incubated with mAbs as described above and used to infect fresh cells for 4 days. P2 supernatants were treated as P1 and P3 supernatants were collected for RNA extraction and subsequent selection step. To select for mAb resistant viruses, 200 µl of the cleared undiluted P3 heterogeneous viral population was incubated with 200 µl mAbs at 2.5 or 10 µg/ml final concentration for 1 hr at 37°C. Mixture was then applied on cells in 800 µl DMEM 2% (1:2 volume) for 3 to 4 days. Viruses were selected one more time and aliquots of passage 5 were used for RNA extraction and sequencing. Virus produced in absence of mAb was collected and treated the same way in parallel to control for appearance of mutations due to cell culture adaptation.

### Spike-pseudotyped vectors production and neutralization assays

HDM-IDT Spike-fixK plasmid (BEI catalogue number NR-52514, obtained from J.D. Bloom, Fred Hutchinson Cancer Research Center) was used as backbone for all the cloning. The cloning of D614G, Alpha, Beta and Delta clones have previously been described (*18*). Pseudoviruses were alternatively produced with the original 2019-nCoV (Cat #100976), Alpha / B.1.1.7 (Cat #101023) and Beta/B.1.351 (Cat #101024) pCAGGS-SARS2-Spike vectors obtained from NIBSC. Omicron ORFs have been cloned with 1kb gBlocks assembly (IDT DNA) followed by In-Fusion cloning in the same plasmid or were generated by gene synthesis with a codon-optimized Spike ORF (Twist Biosciences). Escape mutations have been further introduced by PCR-mediated mutagenesis. Pseudoviruses were produced by co-transfection with pMDL p.RRE, pRSV.Rev and pUltra-Chili-Luc vectors (Addgene) into HEK 293T cells as previously described (*18*).

SARS-CoV-2 pseudotyped VLPs have been produced by co-transfection of HDM-IDT Spike-fixK, CoV-2 N, CoV-2-M-IRES-E and Luc-PS9 plasmids as described in Syed et al (*34*). Briefly, for a 10-cm plate, plasmids CoV-2-N (0.67), CoV-2-M-IRES-E (0.33), HDM-IDTSpike-fixK (0.03) and Luc-PS9 (1.0) at indicated mass ratios for a total of 20 µg of DNA were diluted in 1 ml Opti-MEM (Gibco, ThermoFisher Scientific, USA). Then, 60 µg TransIT- LT1 transfection reagent (Mirus Bio, USA) was added to plasmid dilution to complex the DNA, according to the provider’s instructions. Transfection mixture was incubated for 15 minutes at room temperature and then added dropwise on HEK 293T cells in 10 ml of DMEM containing 10% fetal bovine serum. Media was changed after 18 hours of transfection. VLPs containing media was collected 36 and 48 hours post transfection, pooled, centrifuged 5 minutes at 500 × g and the supernatants filtered using a 0.45 µm syringe filter. Samples were aliquoted and stored at 4°C if used immediately or at -80°C for further analyses.

### RNA genome quantification

Viral RNA was extracted from the supernatants with EZNA viral RNA extraction kit, DNAse-treated when particles were produced by transfection, reverse transcribed with Maxima H Minus cDNA Synthesis Master Mix (ThermoFisher Scientific, USA) as recommended by the manufacturer and the genome quantified by RT-qPCR performed in triplicates using the following primers to detect either the Luciferase gene for VLPs or the RdP gene for viruses: Luc(s): 5- GTG GTG TGC AGC GAG AAT AG -3’; Luc(as): 5- CTG TTC AGC AGC TCG CGC TC -3’; RdP(s): 5-AGC TTG TCA CAC CGT TTC-3’, RdP(as): 5’-AAG CAG TTG TGG CAT CTC-3’.

Absence of DNA contamination was always verified with a control amplification performed in parallel in absence of reverse transcription step.

### Viral escapees sequencing and mapping

Viral RNA was extracted from passage 5 supernatants and deep-sequenced. Sequencing reads were mapped to the SARS-CoV-2 WuhCor1 strain downloaded from the UCSC database, using botwie2 in sensitive mode with read gap penalties 5,1.9. The perbase package (https://github.com/sstadick/perbase) was then used to obtain the nucleotide depth for each base in the genome. Only mutations found in more than 20% of the reads were taken into accounts.

### Infectivity and neutralization assays of pseudotyped particles

In each well of a black 96-well previously coated with poly-L-lysine (0.01% w/v solution, Sigma-Aldrich USA), 50 µl of VLP-containing supernatants were added to 50 µl of cell suspension containing 100 000 receiver cells (HEK293T ACE2/TMPRSS2 cells), in n=8 replicates. 24 hours later, supernatant was removed, then 30 µl of DMEM medium was added with 30 µl of reconstituted luciferase assay buffer (Bright-Glo luciferase assay, Promega, USA) and mixed. Luminescence was measured 5 minutes after using a Hidex Sense microplate reader (Hidex Oy, Finland).

For lentiviral containing supernatants, the protocol is identical except the incubation time is 72 hrs instead of 24 hrs with assays performed as previously described (*17*).

### NHP challenge model for SARS-COV-2 Omicron BA.1 infection

Cynomolgus macaques (*Macaca fascicularis*) originating from Mauritian AAALAC certified breeding centers were used in this study. All animals were housed within IDMIT animal facilities at CEA, Fontenay-aux-Roses under BSL-3 containment when necessary (Animal facility authorization #D92-032-02, Préfecture des Hauts de Seine, France) and in compliance with European Directive 2010/63/EU, the French regulations and the Standards for Human Care and Use of Laboratory Animals, of the Office for Laboratory Animal Welfare (OLAW, assurance number #A5826-01, US). Animals tested negative for Campylobacter, Yersinia, Shigella and Salmonella before being use in the study.

The protocols were approved by the institutional ethical committee “Comité d’Ethique en Expérimentation Animale du Commissariat à l’Energie Atomique et aux Energies Alternatives” (CEtEA #44) under statement number A20-011. The study was authorized by the “Research, Innovation and Education Ministry” under registration number APAFIS#29191- 2021011811505374 v1. All information on the ethics committee is available at https://cache.media.enseignementsup-recherche.gouv.fr/file/utilisation_des_animaux_fins_scientifiques/22/1/comiteethiqueea17_juin2013_257221.pdf.

In the prophylactic protection study, ten female cynomolgus macaques aged 26-27 months at the beginning of the study were randomly assigned between the control and treated groups to evaluate the efficacy of P4J15 LS in protecting from challenge with the SARS-CoV-2 XBB.1.5 virus (NIH/BEI reference: NR-59105; hCoV-19/USA/MD-HP40900/2022). The treated group (n = 6) received one dose at 5 mg/kg of P4J15 LS human IgG1 monoclonal antibody delivered by intravenous slow bolus injection over 3-8 minutes three day prior to challenge, while control animals (n = 4 in parallel and n=2 historical) received no treatment. The two historical control animals were male and infected three weeks before the study with P4J15 LS. All animals were then exposed to a total dose of 10^5^ TCID50 of Omicron XBB.1.5 SARS-CoV-2 virus produced in Vero-ACE2-TMPRSS2 (NIH/BEI reference: NR-59105) via the combination of intranasal and intratracheal routes with sample collection and testing performed as previously described (*35*). Tracheal swabs, nasopharyngeal swabs and bronchoalveolar lavages were performed on all NHPs collected throughout the study to monitor levels of both genomic and subgenomic RNA for the SARS-COV-2 virus as previously described (*36*). All animals and data points were included in the analysis. The NHP sample size was selected based on the large, 1- to 2-log reduction in viral RNA anticipated in the trachea, nasopharyngeal and/or BAL with an effective therapy that can provide statistically significant differences between treated and untreated NHPs. These sample size assumptions were confirmed with the statistical differences observed in viral RNA in viral RNA levels was evaluated using the Mann-Whitney two-sided tests to compare control and treatment groups.

### Hamster challenge model SARS-CoV-2 infection

KU LEUVEN R&D has developed and validated a SARS-CoV-2 Syrian Golden hamster infection model that is suitable for the evaluation of potential antiviral activity of novel antibodies (*37–39*). The SARS-CoV-2 strain used in this study was the Omicron BA.5 (BEI: hCoV-19/USA/COR-22-063113/2022). Infectious virus was isolated by serial passaging on HuH7 and Vero E6 cells (*37*); passage 6 virus was used for the study described here. The titer of the virus stock was determined by end-point dilution on Vero E6 cells by the Reed and Muench method. Live virus-related work was conducted in the high-containment A3 and BSL3+ facilities of the KU Leuven Rega Institute (3CAPS) under licenses AMV 30112018 SBB 219 2018 0892 and AMV 23102017 SBB 219 20170589 according to institutional guidelines.

The hamster infection model of SARS-CoV-2 has been described before (*37, 39*). The animals were acclimated for 4 days prior to study start. Housing conditions and experimental procedures were approved by the ethics committee of animal experimentation of KU Leuven (license P065-2020). Female hamsters of 6-8 weeks old were administered IgG1 isotype control (5 mg/kg), P4J15 LS (5 mg/kg, 1 mg/kg or 0.5 mg/kg) or bebtelovimab (5 mg/kg) by intraperitoneal injection. Two days later, hamsters were anesthetized with ketamine/xylazine/atropine, blood samples were collected, and animals were inoculated intranasally with 2.4×10^6^ median tissue culture infectious dose (TCID50) of SARS-CoV-2 Omicron BA.5 (day 0). Hamsters were monitored for appearance, behavior and weight. Antibody concentrations present in the hamster plasma on day 0 of the study were performed using the Luminex assay described above with Spike trimer coupled beads and using purified P4J15 LS antibody to generate a standard curve. In these studies, no control animals were excluded. In treated groups, animals with undetectable levels of serum antibodies (one hamsters in the 5 mg/kg P4J15 LS group and 2 hamsters in the 1 mg/kg P4J15 LS group) were excluded from the analysis as this indicated a technical failure in the drug administration. At day 4 post infection, hamsters were sacrificed, and lung tissues were homogenized using bead disruption (Precellys) in 350 μl TRK lysis buffer (E.Z.N.A. Total RNA Kit, Omega Bio-tek) and centrifuged (10,000 rpm, 5 min) to pellet the cell debris. RNA was extracted according to the manufacturer’s instructions. Of 50 μl eluate, 4 μl was used as a template in RT-qPCR reactions. RT-qPCR was performed on a LightCycler96 platform (Roche) using the iTaq Universal Probes One-Step RT-qPCR kit (BioRad) with N2 primers and probes targeting the nucleocapsid (*37*). Standards of SARS-CoV-2 cDNA (IDT) were used to express viral genome copies per mg tissue. For end-point virus titrations, lung tissues were homogenized using bead disruption (Precellys) in 350 μl minimal essential medium and centrifuged (10,000 rpm, 5min, 4°C) to pellet the cell debris. To quantify infectious SARS- CoV-2 particles, endpoint titrations were performed on confluent Vero E6 cells in 96- well plates. Viral titers were calculated by the Reed and Muench method using the Lindenbach calculator and were expressed as 50% tissue culture infectious dose (TCID50) per mg tissue. The hamster sample size was selected based on the large, >1-log reduction in viral RNA and infectious virus anticipated in the lung tissue with an effective therapy that can provide statistically significant differences between treated and untreated animals. These sample size assumptions were confirmed in our statistical analysis.

Statistical differences in viral RNA levels and infectivity were evaluated using the Mann-Whitney two-sided tests to compare control and treatment groups.

### Cryo-electron microscopy

Cryo-EM grids were prepared with a Vitrobot Mark IV (ThermoFisher Scientific). Quantifoil R1.2/1.3 Au 400 holey carbon grids were glow-discharged for 90s at 15mA using a GloQube Plus Glow-Discharge System (Quorum, Inc.). 2.0 µl of a 2.1 mg/ml XBB.1 Spike was mixed with 2.0 µl of a 0.28 mg/ml P4J15 Fab fragments (Final 11.1 µM XBB.1 Spike:5.6 µM P4J15 Fab) and 3.0 µl of the fresh complex was applied to the glow-discharged grids, blotted for 4s under blot force 10 at 95% humidity, wait time 10s and 10 °C in the sample chamber, and then the blotted grid was plunge-frozen in liquid nitrogen-cooled liquid ethane.

Grids were transferred in a ThermoFisher Scientific Titan Krios G4 transmission electron microscope, equipped with a Cold-FEG on a Falcon IV detector (Dubochet Center for Imaging, Lausanne) in electron counting mode. Falcon IV gain references were collected just before data collection. Data was collected using TFS EPU v2.12.1 using aberration-free image shift protocol (AFIS), recording 4 micrographs per ice hole. A total of 24 814 micrographs in EER format were recorded at magnification of 165kx, corresponding to the 0.83Å pixel size at the specimen level, with defocus values ranging from -0.6 to -2.0 µm. Exposures were adjusted automatically to 50 e^-^/Å^2^ total dose.

### Cryo-EM image processing

During the data acquisition phase, on-the-fly processing was employed to assess the data quality for screening purposes, utilizing cryoSPARC live v3.3.1 (*40*). Raw stacks were subjected to motion correction without binning, utilizing cryoSPARC’s implementation of motion correction and contrast transfer function estimation (*41*). A total of 6,769,807 particles were automatically template-picked. Following several rounds of 2D classification, 362,329 particles were selected and utilized for ab-initio reconstruction and 3D classifications. Within this dataset, multiple conformers were identified; however, after thorough validation, a subset of 95,910 particles corresponding to a P4J15 fragment bound to an XBB1 trimer was deemed reliable. Homogeneous refinement using the selected particles resulted in a 3D reconstruction at a resolution of 3.01 Å (FSC 0.143) with C1 symmetry. To further enhance the map quality, focused refinement was performed using a soft mask volume encompassing an RBD-up region and its bound Fab. This refinement process yielded a final Coulomb map at 3.85 Å resolution (FSC 0.143) with C1 symmetry (**Supplementary Fig. 4**). The soft mask volumes were manually generated in UCSF ChimeraX (*42*) and the Cryosparc Volume tool. Post-processing polishing was conducted with EMReady (*19*) to improve the map quality and aid in resolving any atom position ambiguities. Finally, the building and refinement steps were exclusively carried out using CryoSPARC maps.

### Cryo-electron microscopy model building

To generate initial models of the P4J15 Fab and XBB1 spike, various approaches were employed. These included utilizing models from the Spike trimer (PDB ID 7QO7), AlphaFold2 (implemented through ColabFold), and ModelAngelo 0.3 (*43*) for sequence-based generation. The cryo-EM maps were fitted with the Spike trimer using UCSF ChimeraX, serving as the starting point for further manual refinement. Manual extension and building of the docked models were carried out using Coot 0.9.8 (*44*). To refine the models, Phenix 1.20 (*45*) was employed. The generated figures depicting the models were prepared using UCSF ChimeraX. The numbering scheme for the full-length Spike models within the global map is based on Omicron numbering. For models containing only the RBD within the local maps, wild-type numbering is utilized. In the case of the P4J15 Fab, the numbering of both the heavy and light chains start from one, beginning with the CH1 and CL domains, respectively. For additional analysis, buried surface area measurements were calculated using ChimeraX. Predictions regarding hydrogen bonds and salt bridges were performed using PDBePISA.

### Statistical analysis

Statistical parameters including the exact value of n, the definition of center, dispersion, and precision measures (Mean or Median ± SEM) and statistical significance are reported in the Figures and Figure Legends. Data were judged to be statistically significant when p < 0.05. In Figures, asterisks denote statistical significance as calculated using the two-tailed non-parametric Mann-Whitney U test for two groups’ comparison or with Kruskal-Wallis tests with Dunn’s multiple-comparison correction. Analyses were performed in GraphPad Prism (GraphPad Software, Inc.) and Microsoft Excel.

### Data availability

All data supporting the findings of this study are available within the paper and in the Source Data. The reconstructed maps of the global Omicron Spike with Fabs bound are available from the EMDB database, C1 symmetry, EMD-17819. The atomic model for the RBD-up with one Fabs bound in the locally refined map is available from the PDB database, PDB-8PQ2. All plasmids made in this study are available upon request to the corresponding authors.

## Author contributions

C.F. designed the strategy for isolating and profiling anti-Spike antibodies, coordinated the research activities, designed Spike variant constructs, analyzed the data, wrote the initial draft and contributed to the editing of the manuscript. P.T. established, performed the experiments with live SARS-CoV-2 virus and designed the Spike protein mutations with the help of C.R., analyzed the results and contributed to the editing of the manuscript. C.R. and V.G. performed Spike cloning and VLP-based experiments. M.L. sets up the SARS-CoV-2 VLPs assay. Y.D., K.L., F.P., L.P., and H.S. coordinated the cryo-EM analysis, analyzed the structural data and contributed to the editing of the manuscript. Other contributed as follows: L.E.-L., performed the B cells sorting, immortalization, binding studies and mAb functional assays; A.F. and J.Ce., cloning of mAb VH and HL, and generation of Omicron BA.5 mutations by site directed mutagenesis; J.Ca., binding studies, production of lentiviruses and pseudoviral assays; F.F. mAb purification, mAb characterization and molecular biology; F.P. coordinated production of recombinant Spike protein and mAb. P.L., Y.L. and R.L. designed the *in vivo* studies, which were executed by C.H, R.M., N. D.-B., F.R., R.A., C.S.F., G.V. and J.N. G.P. and D.T. conceived the study design, analyzed the results and wrote the manuscript.

## Acknowledgements

We thank the Vaccinology and Immunotherapy Centre (VIC) at the Service of Immunology and Allergy of the Lausanne University Hospital for assistance with cell sorting; Laura Junges and Laetitia Bossevot from IDMIT/CEA for qPCR in NHP samples. We thank Isabella Eckerle, Meriem Bekliz and the Virology laboratory of Geneva University Hospital for the Omicron RNA sample and variant isolates collection, the Geneva Genome Center for sequencing and Evarist Planet for the viral sequences mapping. We thank Laurence Durrer, Rosa Schier, Michaël François and Soraya Quinche from the EPFL Protein Production and Structure Core Facility for mammalian cell production and purification of proteins, Alexander Myasnikov, Bertrand Beckert and Sergey Nazarov from the Dubochet Center for Imaging (an EPFL, UNIGE, UNIL initiative) for cryoEM grids preparation and data collection. We thank Julien Lemaitre, Quentin Sconosciuti, Sébastien Langlois, Victor Magneron, Maxime Potier, Jean-Marie Robert, Emma Burban, Eleana Navarre, Flavie Misplon and Tiphaine Bourgès for the NHP experiments; Laura Junges and Kyllian Lheureux for the RT-qPCR. Elodie Guyon, Julien Dinh and Eloise Joffroy for the NHP sample processing; Sylvie Legendre for the transport organization; Frédéric Ducancel, Alicia Pouget and Yann Gorin for the logistics and safety management; Isabelle Mangeot and Salome Piault for help with resources management and Brice Targat and Karl-Stefan Baczowski who contributed to data management. The virus stock used for the NHPs was obtained from the Biodefense, Research Resources, and Translational Research branches of the NIH/NIAID/DMID/OBRRTR/RRS (Program Officer, Clint Florence, Ph.D.). The Infectious Disease Models and Innovative Therapies research infrastructure (IDMIT) is supported by the “Programme Investissements d’Avenir” (PIA), managed by the ANR under references ANR-11-INBS-0008 and ANR-10-EQPX-02-01. We would also like to thank Trudi Veldman and members of the CARE-IMI work package 4 team for helpful discussions. G.P. and R.L. received a grant from the Corona Accelerated R&D in Europe (CARE) project funded by the Innovative Medicines Initiative 2 Joint Undertaking (JU) under grant agreement No 101005077. The JU receives support from the European Union’s Horizon 2020 research and innovation program, the European Federation of Pharmaceutical Industries Associations (EFPIA), the Bill & Melinda Gates Foundation, Global Health Drug Discovery Institute and the University of Dundee. The content of this publication only reflects the author’s view and the JU is not responsible for any use that may be made of the information it contains. G.P. and D.T. received support from the CoVICIS project (grant No. 10146041) funded by the European Union Horizon Europe Program. Additional funding was provided through the Lausanne University Hospital (to G.P.), the Swiss Vaccine Research Institute (to G.P. and NCCR TransCure to H.S.), Swiss National Science Foundation Grants (to G.P.) and through the EPFL COVID fund (to D.T.) and a donation from a private foundation advised by CARIGEST S.A. (to DT and GP).

